# Principal curve approaches for inferring 3D chromatin architecture

**DOI:** 10.1101/2020.06.15.151688

**Authors:** Elena Tuzhilina, Trevor J. Hastie, Mark R. Segal

## Abstract

Three dimensional (3D) genome spatial organization is critical for numerous cellular processes, including transcription, while certain conformation-driven structural alterations are frequently oncogenic. Genome architecture had been notoriously difficult to elucidate, but the advent of the suite of chromatin conformation capture assays, notably Hi-C, has transformed understanding of chromatin structure and provided downstream biological insights. Although many findings have flowed from direct analysis of the pairwise proximity data produced by these assays, there is added value in generating corresponding 3D reconstructions deriving from superposing genomic features on the reconstruction. Accordingly, many methods for inferring 3D architecture from proximity d hyperrefata have been advanced. However, none of these approaches exploit the fact that single chromosome solutions constitute a one dimensional (1D) curve in 3D. Rather, this aspect has either been addressed by imposition of constraints, which is both computationally burdensome and cell type specific, or ignored with contiguity imposed after the fact. Here we target finding a 1D curve by extending principal curve methodology to the metric scaling problem. We illustrate how this approach yields a sequence of candidate solutions, indexed by an underlying smoothness or degrees-of-freedom parameter, and propose methods for selection from this sequence. We apply the methodology to Hi-C data obtained on IMR90 cells and so are positioned to evaluate reconstruction accuracy by referencing orthogonal imaging data. The results indicate the utility and reproducibility of our principal curve approach in the face of underlying structural variation.

## 1 Introduction

The three-dimensional (3D) configuration of chromosomes within the eukaryote nucleus is important for several cellular functions, including gene expression regulation, and has also been linked to translocation events and cancer driving gene fusions (Mitelman *and others*, 2007). While direct visualization of 3D architecture has improved (see Section 2.9), imaging challenges pertaining to chromatin compaction and dynamics persist. However, the ability to *infer* chromatin architectures at increasing resolution has been enabled by chromosome conformation capture (3C) assays (Dekker *and others*, 2002). In particular, when coupled with next generation sequencing, such Hi-C methods (Lieberman-Aiden and *others*, 2009; Duan *and others*, 2010) yield an inventory of pairwise, genome-wide chromatin interactions, or contacts. In turn, the contact data form the basis for *reconstructing* 3D configurations (Zhang *and others*, 2013; Varoquaux *and others*, 2014; Ay *and others*, 2014; Zou *and others*, 2016; Rieber and Mahony, 2017). While many novel conformational-related findings have flowed from direct analysis of contact level data, added value of performing downstream analysis based on attendant 3D reconstructions has been demonstrated. These benefits derive from the ability to superpose genomic features on the reconstruction. Examples include co-localization of genomic landmarks such as early replication origins in yeast (Witten and Noble, 2012; Capurso and Segal, 2014), gene expression gradients in relation to telomeric distance and co-localization of virulence genes in the malaria parasite (Ay *and others*, 2014), the impact of spatial organization on double strand break repair (Lee *and others*, 2016), and elucidation of ‘3D hotspots’ corresponding to (say) overlaid ChIP-Seq transcription factor extremes which can reveal novel regulatory interactions (Capurso *and others*, 2016).

The contact or interaction matrices resulting from Hi-C assays, which are typically performed on bulk cell populations, are depicted as heatmaps, which record the frequency with which pairs of binned genomic loci are cross-linked, reflecting spatial proximity of the respective loci bins within the nucleus. A common first step toward 3D reconstruction is the conversion of contact frequencies into *distances*, typically assuming inverse power-law relationships (Varoquaux *and others*, 2014; Ay *and others*, 2014; Shavit *and others*, 2014; Rieber and Mahony, 2017), from which 3D chromatin architecture can be obtained via versions of the multi-dimensional scaling (MDS) paradigm. In response to (i) the bulk cell population underpinnings of contact data, (ii) computational challenges posed by the dimensionality of the MDS reconstruction problem as governed by bin extent, and (iii) accommodating biological considerations, several competing reconstruction algorithms have been advanced. However, none of these take advantage of the fact that the 3D solution for individual chromosomes corresponds to a one-dimensional (1D) curve in 3-space. Rather, this aspect has been addressed by imposition of constraints (Duan *and others*, 2010; Ay *and others*, 2014; Stevens *and others*, 2017), which are cell type specific and require prescription of constraint parameters. These parameters can be difficult to specify and their inclusion substantially increases the computational burden. Other approaches (Zhang *and others*, 2013; Park and Lin, 2017; Rieber and Mahony, 2017) do not formally incorporate contiguity but impose it post hoc, creating chromatin reconstructions by “connecting the dots” of the 3D solution according to the ordering of corresponding genomic bins.

Here we directly target chromosome reconstruction by finding a 1D curve approximation to the contact matrix via extending principal curve methodology (Hastie and Stuetzle, 1989) to the metric scaling problem. After reviewing problem formulation and current reconstruction techniques in Section 2.1, we develop two building blocks, *Principal Curve Metric Scaling* (PCMS; Sections 2.2, 2.3) and *Weighted PCMS* (WPCMS; Sections 2.4, 2.5), that enable our novel *Poisson Metric Scaling* (PoisMS; Sections 2.6, 2.7) approach. Strategies for selecting a specific reconstruction from a degrees-of-freedom indexed series of solutions are described in Section 2.8. Methods for appraising the accuracy of candidate reconstructions using orthogonal imaging data are outlined in Section 2.9. Results from applying the methodology to Hi-C data from IMR90 cells are presented in Section 3, while the Discussion indicates directions for future work.

## 2 Methods

### 2.1 Existing approaches to 3D chromatin reconstruction from Hi-C assays

Our focus is on reconstruction of *individual* chromosomes; whole genome architecture can follow by appropriately positioning these solutions (Segal and Bengtsson, 2015; Rieber and Mahony, 2017). As is standard, we disregard complexities deriving from chromosome pairing arising in diploid cells (which can be disentangled at high resolutions (Rao *and others*, 2014)) and address issues surrounding bulk cell experiments and inter-cell variation in the Discussion.

The result of a Hi-C experiment is the *contact map*, a symmetric matrix 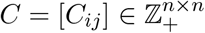 of contact counts between *n* (binned) genomic loci *i,j* on a genome-wide basis; Figure 1 provides an example. We defer questions surrounding contact matrix normalization. This matrix can be exceedingly sparse, even after binning. The 3D chromatin reconstruction problem is to use the contact matrix *C* to obtain a 3D point configuration 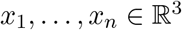 corresponding to the spatial coordinates of loci 1,…, *n* respectively; Figure 2 gives an illustration.

**Figure 1:**
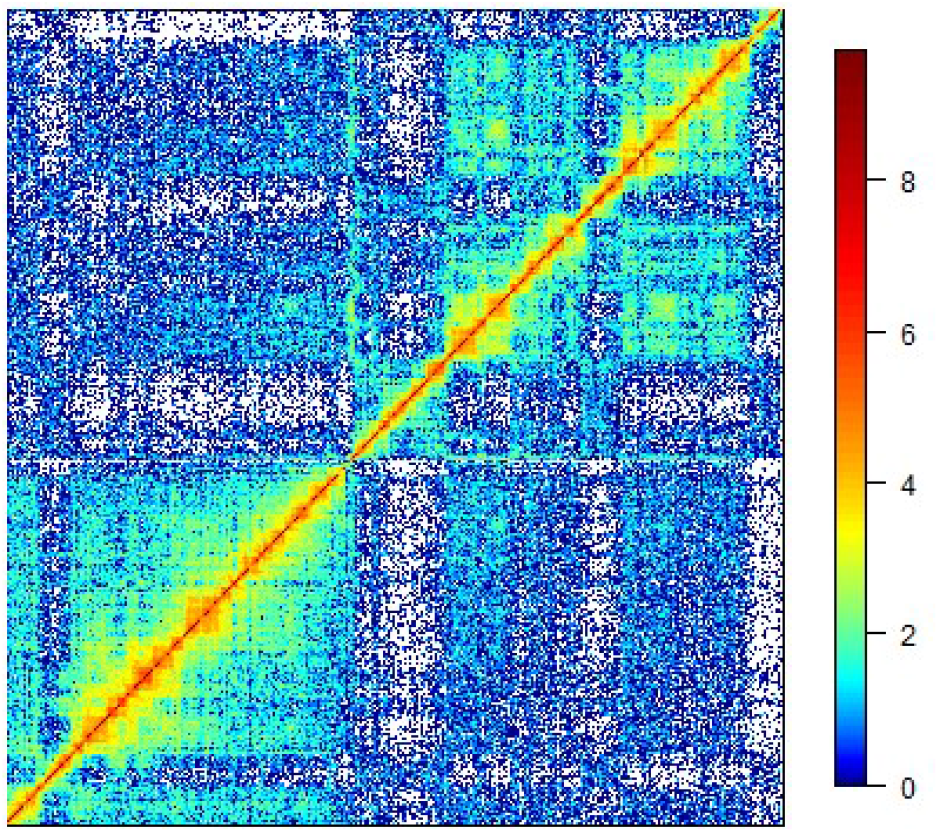
Log-transformed contact matrix log(*C*). White color corresponds to *C_ij_* = 0 or, equivalently, log(*C_ij_*) = −∞.

Many approaches have been proposed to tackle this problem with broad distinction between optimization and modelbased methods (Varoquaux *and others*, 2014; Rieber and Mahony, 2017). A common first step is conversion of the contact matrix into a distance matrix 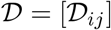 (Duan *and others*, 2010; Varoquaux *and others*, 2014; Ay *and others*, 2014; Shavit *and others*, 2014), followed by solving the *multi-dimensional scaling* (MDS; Hastie *and others*, 2009) problem: position points (corresponding to genomic loci) in 3D so that the resultant interpoint distances best conform to the distance matrix.

A variety of methods have also been used for transforming contacts to distances. At one extreme, in terms of imposing biological assumptions, are methods that relate observed intra-chromosomal contacts to genomic distances and then ascribe *physical* distances based on organism specific findings on chromatin packing (Duan *and others*, 2010) or relationships between genomic and physical distances for crumpled polymers (Ay *and others*, 2014). Such distances inform the subsequent optimization step as they permit incorporation of known biological constraints that can be expressed in terms of physical separation. Importantly, these constraints include prescriptions on the 3D separation between contiguous genomic bins. It is by this means that obtaining a 1D curve is indirectly facilitated. However, obtaining physical distances requires both strong assumptions and organism specific data (Fudenberg and Mirny, 2012). More broadly, a number of approaches (Zhang *and others*, 2013; Varoquaux *and others*, 2014; Zou *and others*, 2016; Rieber and Mahony, 2017) utilize power law transfer functions to map contacts to (non-physical) distances 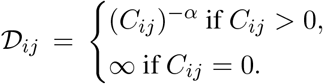. Adoption of the power law derives from empirical and theoretical work but again constitutes a strong assumption (Fudenberg and Mirny, 2012).

Once we have a distance matrix 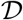, optimization approaches seek a 3D configuration *x*_1_,…, *x_n_* that best fits 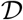 according to an MDS criterion. If ‖·‖ designates the Euclidean norm, then an example of MDS loss incorporating weights and penalty (Zhang *and others*, 2013) is

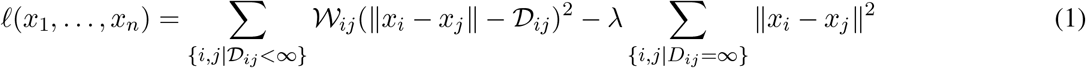

with the corresponding optimization problem

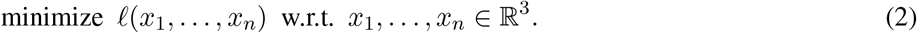

Here common choices for the weights 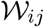 include 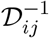 (Zhang *and others*, 2013) and 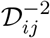 (Varoquaux *and others*, 2014), these being analogous to precision weighting since large *C_ij_* (small 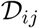) are more accurately measured. Similarly, the penalty (second) term maximizes the pairwise distances for loci bins with *C_ij_* = 0 under the presumption that such loci should not be too close.

It is worth noting that (1), and related criteria, correspond to a nonconvex, nonlinear optimization problem that is NP hard and while various devices have been employed to mitigate the computational burden (e.g., Zhang *and others*, 2013), computational concerns, particularly for high resolution (many loci bins) problems, remain forefront.

Probabilistic methods model the contact counts with an optimization goal of maximizing the corresponding log-likelihood. In particular, Poisson models, *C_ij_* ~ *Pois*(λ_*ij*_), are widely used (Varoquaux *and others*, 2014; Zou *and others*, 2016; Park and Lin, 2017), where λ_*ij*_ = λ_*ij*_(*x*_1_,…, *x_n_*) is a function of the genomic loci spatial coordinates *x*_1_,…, *x_n_*. For example, Rosenthal *and others* (2019) prescribe exponential dependence between the Poisson rate parameter and inter-loci distances: λ_*ij*_ = *β*‖*x_i_* – *x_j_*‖^*α*^ for some *α* < 0, a framework we slightly modify in Section 2.6.

All existing approaches implicitly represent chromatin as a polygonal chain. Constraints on the geometrical structure of the polygonal chain can be imposed via penalties on edge lengths and angles between successive edges, with even quaternion-based formulations employed (Caudai *and others*, 2015). Rosenthal *and others* (2019) utilize penalties to control smoothness of the resulting conformations. However, despite imparting targeted properties to the resulting reconstruction, such penalty-based approaches increase the complexity of the objective, its gradient and Hessian, both slowing and limiting, especially with respect to resolution, associated algorithms.

Here we develop a suite of novel approaches that directly model chromatin configuration as a 1D curve in 3D. Our primary method, *Poisson Metric Scaling* (PoisMS), is based on a Poisson model for contact counts and provides an efficient means for obtaining smooth 1D reconstructions, that combines advantages of both MDS and probabilistic models. This technique utilizes two building blocks of intrinsic interest. First, we introduce the *Principal Curve Metric Scaling* (PCMS) approach that features an optimization problem inspired by MDS and stated in terms of inner products. This problem admits a simple solution obtained via the singular value decomposition. Next, we develop *Weighted PCMS* (WPCMS), a weighted version of PCMS that, importantly, models distances rather than inner products and further permits control over the influence of particular elements of the contact matrix on the resulting reconstruction. This technique requires an iterative algorithm that uses PCMS as the core component. Finally, WPCMS in turn can be used in conjunction with projected gradient descent to solve a second order approximation of the Poisson log-likelihood, yielding our PoisMS algorithm.

### 2.2 PCMS: metric scaling with a smooth curve constraint

The PCMS technique is based on classical MDS. Given a symmetric matrix *Z*, PCMS treats it as a similarity matrix and approximates it by an inner product matrix (Buja *and others*, 2008). In particular, *Z* can correspond to the contact matrix after conversion to a distance matrix followed by double centering, the standard MDS device that turns (Euclidean) squared distances into inner products. We illustrate this approach in the Supplement (S2) with distances obtained via power law transformation. However, while it is thereby possible to use PCMS as a standalone reconstruction tool, we seek methods that avoid having to convert contacts to distances. So, here we develop PCMS with a view to utilizing it as a building block of our PoisMS technique.

Let 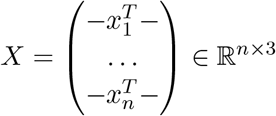 be the matrix of genomic loci coordinates and let *S*(*X*) = *XX^T^* refer to the inner product matrix of the reconstruction *X*. If ‖·‖_*F*_ denotes the Frobenius norm, then the goal is to minimize the *Strain* objective:

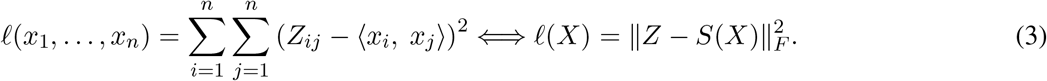

Instead of adding a smoothness penalty to the objective, we impose an additional constraint:

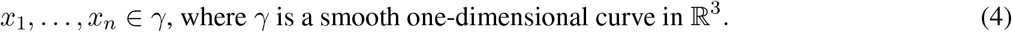

This constraint will serve to capture the inherent contiguity of chromatin. We model the curve *γ* by a cubic spline with *k* degrees-of-freedom as follows (Hastie *and others*, 2009). Suppose *h*_1_(*t*),…, *h_k_*(*t*) are cubic spline basis functions in 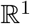 then

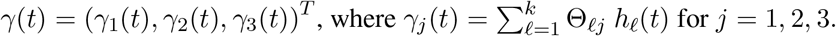

Let *t_i_* index the genomic locus of *x_i_* in the parameter space of *γ*, i.e. *x_i_* = *γ*(*t_i_*), and 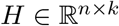 be the matrix of spline basis evaluations at *t_i_*, i.e. *H_iℓ_* = *h_ℓ_*(*t_i_*). Since binning typically results in evenly spaced genomic loci it is convenient to set *t*_1_ = 1, *t*_2_ =2,…, *t_n_* = *n*, although irregular spacing is readily handled. So, the constraint (4) can be written as 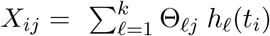, or equivalently, in matrix form as *X* = *H*Θ leading to the optimization problem

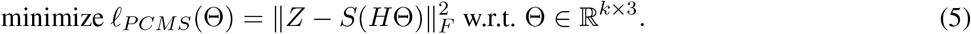

Hereafter we denote the corresponding solution by 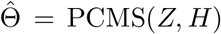, the resulting chromatin reconstruction by 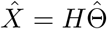 and the approximation of the original matrix *Z* as 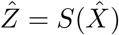.

### 2.3 PCMS solution via eigen-decomposition

Note that the parameter Θ in the PCMS problem (5) is unconstrained. Since Θ is defined up to a multiplication by a full-rank matrix, we can assume *H* to be a matrix with orthogonal columns. To find the PCMS solution the following lemma, proved in Supplement Section S1, is useful.

#### Lemma 2.1

If 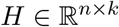 is a matrix with orthogonal columns, i.e. *H^T^H* = *I*, then problem (5) is equivalent to

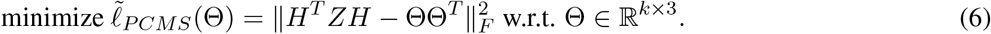

Minimizing 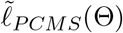 can be interpreted as approximating the matrix *H^T^ZH* by a positive semi-definite rank 3 matrix ΘΘ^*T*^. Assuming that the symmetric matrix *H^T^ ZH* has at least three positive eigenvalues the solution can be found via eigen-decomposition of *H^T^ ZH*: let *H^T^ ZH* = *Q*Λ*Q^T^* for orthogonal *Q* and diagonal Λ = diag(λ_1_,…, λ_*n*_) with λ_1_ ≥ λ_2_ ≥ … ≥ λ_*n*_, then

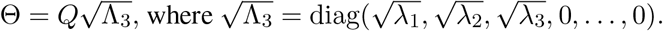

The computational efficiency of PCMS derives from the fact that it relies on eigen-decomposition of a small *k* × *k* matrix, requiring only *O*(*k*^3^) additional operations.

### 2.4 WPCMS: a distance-based model for chromatin reconstruction

As indicated, direct application of PCMS to Hi-C data is limited by the need to convert contact counts to distances and then (via double centering) to inner products since such conversion can be problematic. Even simplistic approaches, based on power law transformation, prescribe a value for the index parameter, failing to accommodate dependence of the index on influencing factors such as cell type, chromosome, organism and resolution. Moreover, the double centering trick requires that resultant distances be Euclidean.

Accordingly, we develop a distance-based version of PCMS, wherein the symmetric matrix *Z* contains pairwise squared distances, as opposed to inner products. Additional flexibility is facilitated by introducing weights to the problem setup, which permits control over the impact of particular elements *Z_ij_* on the reconstruction, for example to counteract diagonal dominance (Yang *and others*, 2017). Although the resulting technique, *Weighted* PCMS (WPCMS), can again be used as a standalone reconstruction tool (Supplement Section S4), akin to PCMS its primary purpose is as component of the PoisMS approach.

We introduce a matrix of weights *W* ∈ [0, 1]^*n*×*n*^, denote by *D*(*X*) the matrix of pairwise distances between genomic loci and consider the following loss function

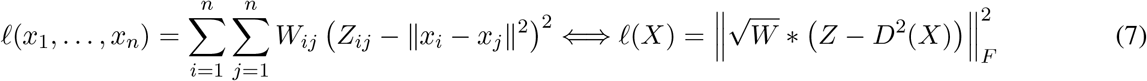

where * refers to the Hadamard (element-wise) product and matrix squaring is also element-wise. The WPCMS problem can be stated as follows:

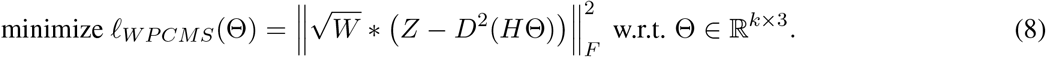

The corresponding solution and reconstruction are denoted by 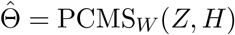 and 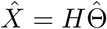, respectively, along with the corresponding approximation 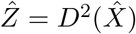 of matrix *Z*.

### 2.5 Iterative algorithm for solving the WPCMS problem

Problem (8) can be elegantly solved using *projected gradient descent* (PGD) (Hastie *and others*, 2015), broadly used to solve constrained optimization problems. We first exploit the fact that the matrix of squared distances can be rewritten in terms of the inner product matrix:

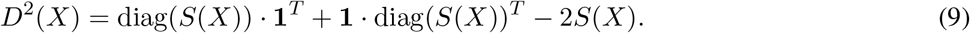

Here 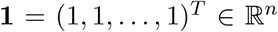 and diag(*S*(*X*)) = (‖*x*_1_‖^2^, ‖*x*_2_‖^2^,…, ‖*x_n_*‖^2^)^*T*^ is the diagonal of the inner product matrix. So (8) can be restated in terms of inner products:

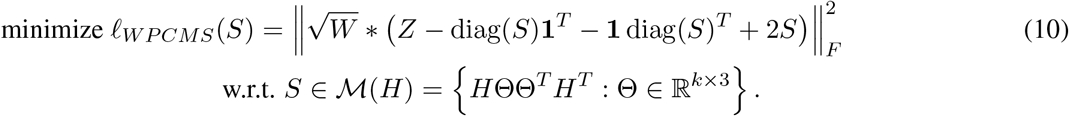

The PGD procedure alternates the following two steps:

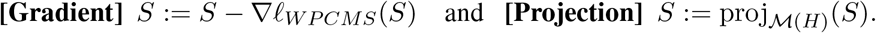

Here 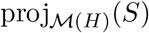 denotes the projection of matrix *S* onto the matrix manifold 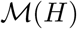. The **[Gradient]** step makes recourse to the following Lemma, proved in Supplement Section S3.

#### Lemma 2.2

Let *D*^2^ denote the matrix of squared distances corresponding to inner product matrix *S* (as in 9). If *G* = *W* * (*Z* − *D*^2^) and *G*_+_ = diag(*G* · **1**) is the diagonal matrix containing row sums of *G* on the diagonal, then up to a scaling factor ∇_*ℓWPCMS*_(*S*) = *G* – *G*_+_.

Next, note that the **[Projection]** step requires solving the optimization problem

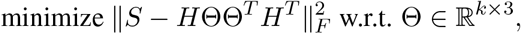

which is easily done using PCMS. Thus, we end up with the following PGD procedure:

1. **[Initialize]** Generate random 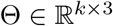, set the reconstruction *X* = *H*Θ.
2. *Repeat until convergence*:

2.1 **[SDG]** Calculate the current guess for the inner product matrix *S* = *XX^T^* and use it to compute the matrix of squared distances *D*^2^ = diag(*S*) · **1**^*T*^ + **1** · diag(*S*)^*T*^ – 2*S*. Then compute *G* = *W* * (*Z* – *D*^2^) as well as *G*_+_ = diag(*G* · **1**).
2.2 **[Gradient]** Update matrix of inner products *S*: = *S* – (*G* – *G*_+_).
2.3 **[Projection]** Update spline coefficients using **PCMS** Θ:= PCMS(*S, H*), then update the reconstruction *X* = *H*Θ.

Convergence is assessed via the stopping criterion 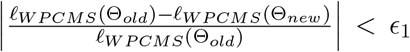, where *ϵ*_1_ is a pre-chosen accuracy rate, and Θ_*old*_ and Θ_*new*_ are Θ values calculated at the previous and current iterations respectively. Details and extensions of WPCMS are provided in the Supplement.

### 2.6 PoisMS: Poisson model for contact counts

We now develop our primary approach, Poisson Metric Scaling (PoisMS), using WPCMS as a building block. We define a probabilistic model for contact counts based on natural and previously adopted assumptions: Poisson distributed counts *C_ij_* with dependence of the Poisson mean on chromatin 3D structure, specifically on pairwise (squared) distances between genomic loci:

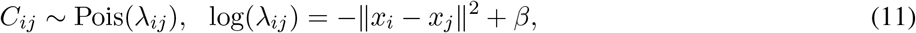

with 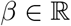 an intercept parameter. The negative log-likelihood objective is

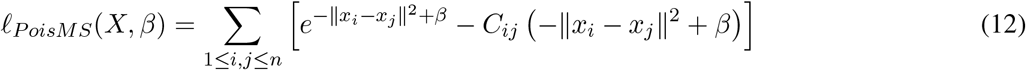

and the MLE optimization problem under the smooth curve constraint (4) is

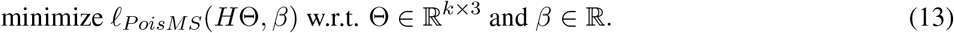

In (11) we use squared distance rather than distance, reflecting criterion (1) and conferring computational convenience. We denote the corresponding matrix of spline coefficients by 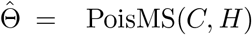 and the resulting chromatin reconstruction by 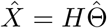.

### 2.7 Iterative algorithm for solving PoisMS problem

A virtue of the Poisson model is that the second order Taylor approximation (SOA) of the negative log-likelihood (12) is simply the weighted Frobenius norm. Further, it is well known that the optimal value of this SOA amounts to one step of the Newton method for optimizing the original loss function. We use these facts to develop an iterative algorithm based on the WPCMS technique, which is equivalent to a projected Newton Method.

First, we review the SOA of the negative Poisson log-likelihood in the univariate case. Suppose *c* ~ Pois(λ). The negative log likelihood *ℓ*(λ) = λ – *c*log λ can be reparametrized in terms of the natural parameter *η* = log(λ) leading to *ℓ*(*η*) = *e^η^* – *cη*. Then the SOA of the reparametrized negative log-likelihood at some point *η*_0_ = log λ_0_, up to scaling and shifting by a constant, is:

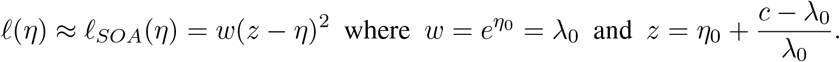

The multivariate version is as follows. Suppose 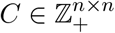 where *C_ij_* ~ Pois(λ_*ij*_) and *η_ij_* = log(λ_*ij*_). Let the respective matrices of Poisson and natural parameters be 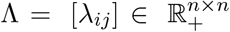 and 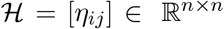. Then the SOA of the negative log-likelihood at some point 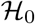, up to scale and shift constants, is

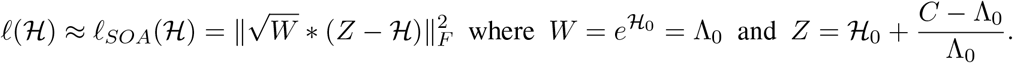

Here * is the Hadamard (element-wise) product, with matrix exponentiation and division also being interpreted as element-wise operations.

Recall that in the Poisson model (11) the natural parameter depends linearly on the matrix of genomic loci pairwise distances: 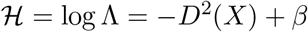. So, the SOA can be rewritten as

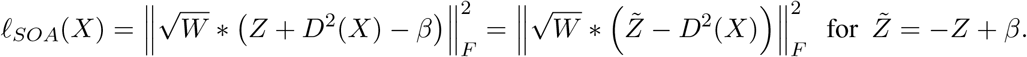

Suppose that the current reconstruction guess is *X*_0_ with corresponding natural parameter value 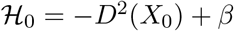. Then we have the following approximation of the Poisson loss (12) at point 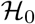 again up to scaling and shifting by a constant:

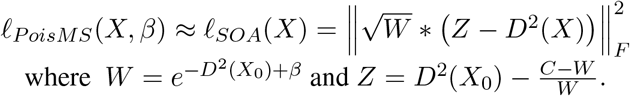

Thus, under the smooth curve constraint *X* = *H*Θ, the loss function *ℓ_SOA_*(*X*) coincides with the WPCMS loss (8) and we obtain a nice application of the WPCMS algorithm, with the solution to the second order approximation of problem (13):

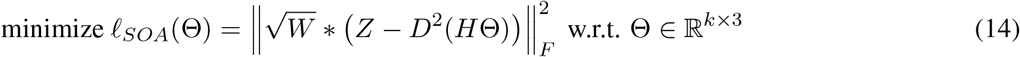

being exactly Θ = PCMS_*W*_(*Z, H*). This observation can be applied to simplify computations for the Poisson model and underlies our PoisMS algorithm.

The last step of our PoisMS algorithm is to update *β* according to the current guess of Θ. This can be done by optimizing the negative log-likelihood with respect to *β*. All together this leads to the following algorithm that repeatedly approximates the Poisson objective at current guess Θ by a quadratic function and shifts Θ towards the global minimum of this quadratic approximation:

1. **[Initialize]** Generate random 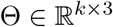, set the reconstruction *X* = *H*Θ.
2. *Repeat until convergence*:

2.1 **[Update *β*]** Update the intercept 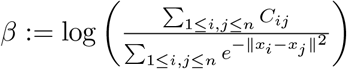.
2.2 **[SOA]** Calculate SOA matrices *W* = *e*^−*D*^2^(*X*) + *β*^ and 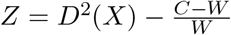.
2.3 **[WPCMS]** Update the spline coefficients using WPCMS approach Θ:= PCMS_*W*_(*Z, H*), then update the reconstruction *X* = *H*Θ.

The stopping rule for the PoisMS algorithm is similar to WPCMS: for some fixed accuracy rate *ϵ*_2_ we check if the updated (Θ_*new*_, *β_new_*) meets the criteria 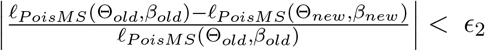 after each iteration of steps 2.1–2.3.

The non-convexity of the PoisMS criteria (13) implies that initialization can impact the resulting reconstruction. In the Supplement we discuss use of WPCMS to provide a warm start for the PoisMS algorithm, as well as algorithmic extensions and computational complexity.

### 2.8 Determination of principal curve degrees-of-freedom

The main hyperparameter of the PoisMS approach is the spline degrees-of-freedom *df* (spline basis size), which controls the smoothness of the resulting reconstruction. To determine the optimal value, for each *df* we create the spline basis matrix *H_df_*, find the corresponding solution 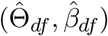 and the resulting reconstruction 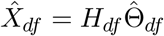. We measure the error rate by the normalized Poisson deviance, i.e

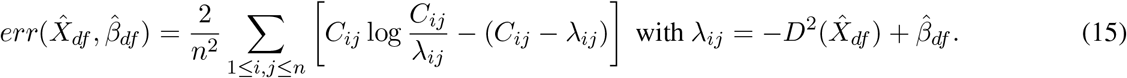

Initially, we tried cross-validation to find the optimal value of *df*, as is common for smoothing (penalty) parameter determination. However, the complex and structural dependencies that characterize contact matrices made this approach problematic. As an alternative we adopted an approach based on identifying the “elbow” that is prototypic in graphs of resubstitution error, here 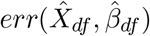, versus model complexity, here *df*. The logic as to why this change point constitutes a basis for model complexity determination is described in Breiman *and others* (1984) in terms of bias-variance tradeoff. Elbow identification is also used for determining appropriate numbers of principal components (Jolliffe, 2002) and clusters (Hastie *and others*, 2009), as well as dimension in MDS (Kruskal and Wish, 1978) and non-negative matrix factorization (NMF; see Hutchins *and others*, 2008) problems.

### 2.9 Accuracy assessment via multiplex FISH

While the prescription in Section 2.8 provides a means for selecting a particular PoisMS model it does not address the accuracy of the chosen model. The absence of gold standards makes such assessment challenging. In comparing competing 3D genome reconstructions several authors have appealed to simulation (Zhang *and others*, 2013; Varoquaux *and others*, 2014; Zou *and others*, 2016; Park and Lin, 2017), however, real data referents are preferable. To that end, many of the same reconstruction algorithm developers have made recourse to fluorescence in situ hybridization (FISH) imaging as a basis for gauging accuracy. This proceeds by comparing distances between imaged probes with corresponding reconstruction-based distances. But such methods are necessarily limited by the sparse number of probes (~ 2 – 6; see Lieberman-Aiden *and others*, 2009; Shavit *and others*, 2014; Park and Lin, 2017) and the modest resolution thereof, many straddling over 1 megabase. The recent advent of *multiplex* FISH (Wang *and others*, 2016) transforms 3D genome reconstruction accuracy evaluation by providing an order of magnitude more probes and hence two orders of magnitude more inter-probe distances than conventional FISH. Moreover, the probes are at higher resolution and centered at topologically associated domains (TADs; see Dixon *and others*, 2012). We use this imaging data, along with companion accuracy assessment approaches (Segal and Bengtsson, 2018) to evaluate our PoisMS reconstructions.

The image-based 3D genomic coordinates furnished from multiplex FISH serve to define the gold standard by which we assess reconstructions. The existence of numerous multiplex FISH replicates is crucial for this task and three steps are necessary to effect such evaluation.

#### Obtaining the gold standard

Given *N* multiplex FISH replicates denote the matrix of the spatial coordinates for replicate *i* ∈ {1,…, *N*} by 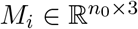 where *n*_0_ denotes the number of distinct multiplex FISH loci (probes) over all replicates. We start by defining the *medoid replicate*. For a pair of 3D conformations *X*_1_, *X*_2_ ∈ **R**^*n*_0_×3^ denote the number of observed loci by *n*(*X*_1_, *X*_2_) and suppose *d_proc_*(*X*_1_, *X*_2_) is the squared Procrustes distance from *X*_2_ to *X*_1_ following alignment allowing translation, rotation and scaling (Hastie *and others*, 2009). Then the dissimilarity between *X*_1_ and *X*_2_ is defined by

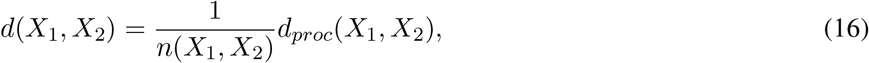

using asymmetric (scaling and rotation transforms applied to *X*_2_ only) Procrustes distance. This measure of agreement between two reconstructions coincides with mean squared deviation (see, for example, Segal and Bengtsson (2018)).

We next define the *medoid* replicate as the replicate whose (weighted) average dissimilarity to the other replicates is minimal:

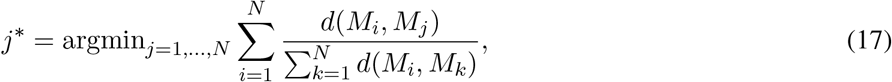

with weights 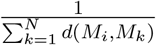 chosen to adjust for different scales of the multiplex FISH replicates. Next, let 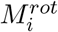 be the Procrustes alignment of *M_i_* to the medoid *M_j*_*. The *average Procrustes conformation* 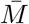, defined as the locus-wise average of the 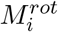, then serves as a gold standard. Our application of Procrustes alignment prior to this (noise reducing) averaging accommodates translation, rotation, and scaling differences between replicate conformations.

#### Computing the reference distribution

Treating the average Procrustes conformation 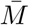 as our gold standard we obtain a reference distribution by measuring the dissimilarity between it and the multiplex FISH replicates: 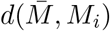. The resulting empirical distribution captures experimental variation around the gold standard. A fine point is that this distribution will exhibit reduced dispersion compared to its target population quantity owing to data re-use since *M_i_* contributes to 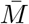. While this concern could be mitigated by employing leave-one-out techniques the large number of available replicates (> 110) renders this unnecessary (Segal and Bengtsson, 2018).

#### Evaluating chromatin reconstructions

To evaluate reconstructions resulting from the PoisMS approach we first need to align the reconstruction with the gold standard. This may involve preliminary coarsening of one or other coordinate sets to yield comparable resolution. Here, the genomic coordinate ranges for each multiplex FISH probe are coarser than the Hi-C bins used in our reconstructions. So we calculate the average of the reconstruction coordinates falling in the corresponding multiplex FISH bins to obtain a lower resolution reconstruction 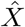 of the same dimension as 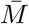. To quantify how close this reconstruction is to the gold standard 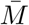 we again measure dissimilarity following alignment 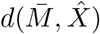 Interpretations of this quantity in the context of the reference distribution are presented in the Results section.

### 2.10 A contrasting reconstruction algorithm: HSA

To compare our PoisMS solution with an alternate reconstruction algorithm we make recourse to HSA (Zou *and others*, 2016). This technique provides an interesting contrast in that it employs a similar Poisson formulation to (12) but instead of contiguity being captured via principal curves per (4), it is indirectly imparted by constraints that induce dependencies on a hidden Gaussian Markov chain over the solution coordinates. Obtaining these spatial coordinates is achieved via simulated annealing with further smoothness effected via distance-based penalization.

HSA has performed well in some benchmarking studies and features several compelling attributes including (i) simultaneously handling multiple data tracks allowing for integration of replicate contact maps and (ii) adaptively estimating the power-law index whereby contacts are transformed to distances as previously emphasized. Nonetheless, in contrast to PoisMS, HSA incurs a substantial compute and memory burden, and questions surrounding robustness have been raised (Rieber and Mahony, 2017).

To compare PoisMS performance with HSA we use the approach described in Section 2.9. Having obtained a HSA reconstruction we measure the dissimilarity between the reconstruction and the gold standard. The quantity so obtained is interpreted in the context of the attendant reference distribution (see Section 3.3 and Supplement Section S10).

## 3 Results

### 3.1 Chromosome reconstructions

We present PoisMS reconstructions for IMR90 cell chromosome 20 at 100kb resolution for which multiplex FISH and Hi-C data acquisition and processing has been previously described (Segal and Bengtsson, 2018). Results for chromosome 21 are presented in the Supplement.

In Figure 1 we present the heatmap for log(*C*). The resulting PoisMS reconstructions 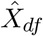 along with the Poisson parameter matrix 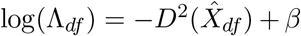, that can be viewed as an approximation of log(*C*), are presented in Figure 2 for a series of degrees-of-freedom values.

**Figure 2:**
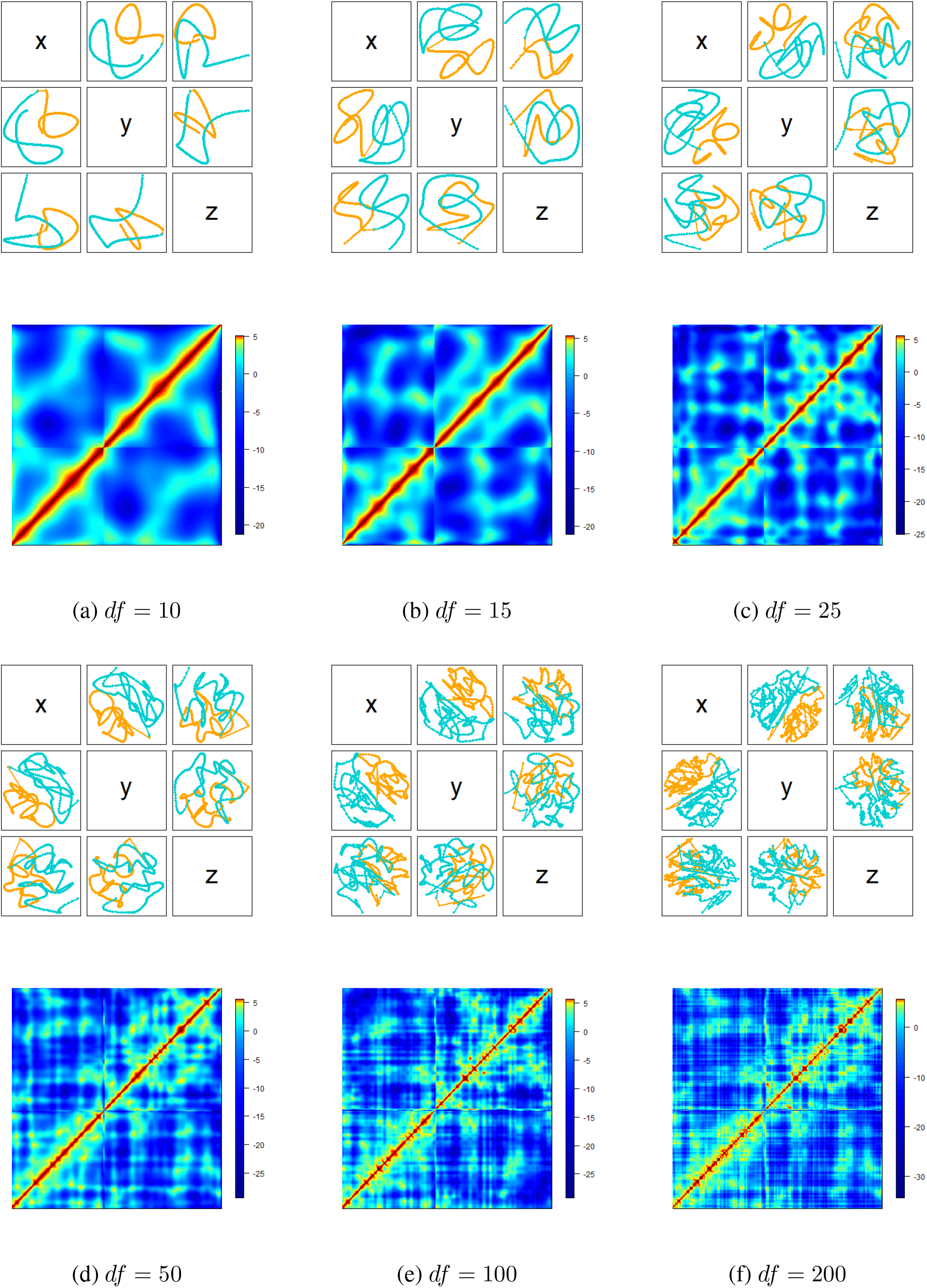
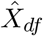, the projections of the resulting reconstruction, with colors (orange, teal) distinguishing chromosome arms, and 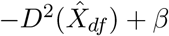, the approximation of log(*C*), obtained via PoisMS for differing degrees-of-freedom values *df*.

### 3.2 Determining degrees-of-freedom

The graph of error rate 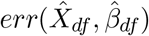 vs *df* reveals rapidly decreasing error rates up to *df* = 30 with subsequent gradual decline (Figure 3). The optimal *df* according to the elbow heuristic, obtained using the **R** package `segmented` (Muggeo, 2008), is *df* = 25, also shown in Figure 3.

**Figure 3:**
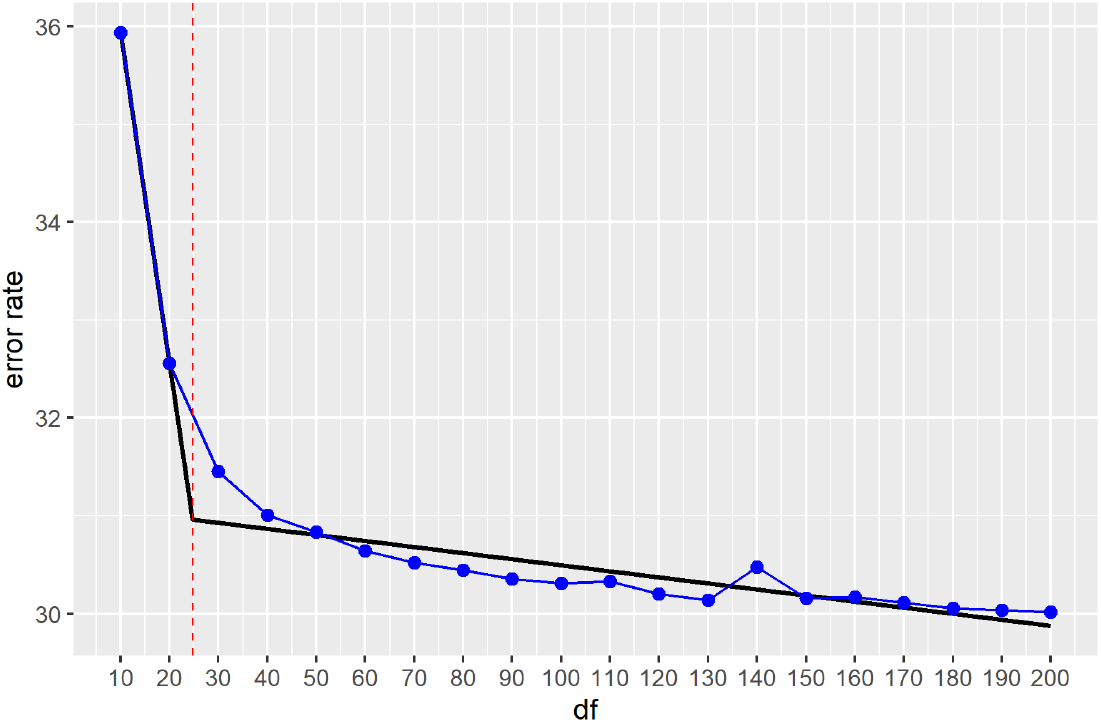
Error rate 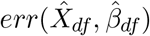 vs. degrees-of-freedom *df* plot for the PoisMS approach. The segmented regression is given by the piecewise linear fit (black) with the degrees-of-freedom selected via kink estimation indicated by the red vertical line and segmentation change point corresponding to *df* = 25.

### 3.3 Evaluating reconstructions via the multiplex FISH referent

Procrustes alignment of 3D conformations, and calculation of the corresponding distances *d_proc_*(·, ·), was performed using the R package `vegan` (Oksanen *and others*, 2016). We obtain the multiplex FISH medoid conformation based on the smallest row sum (17) of the dissimilarity matrix of normalized Procrustes distances (16) as described above. The 111 multiplex FISH replicate conformations are then aligned to the medoid as a prelude to calculating the average Procrustes conformation – our gold standard. Figure 4 shows the histogram of dissimilarities between multiplex FISH replicates and our derived gold standard that constitutes the reference distribution. We position the PoisMS reconstruction dissimilarities therein corresponding to the indicated series of degrees-of-freedom values. HSA reconstruction dissimilarity values are also included.

**Figure 4:**
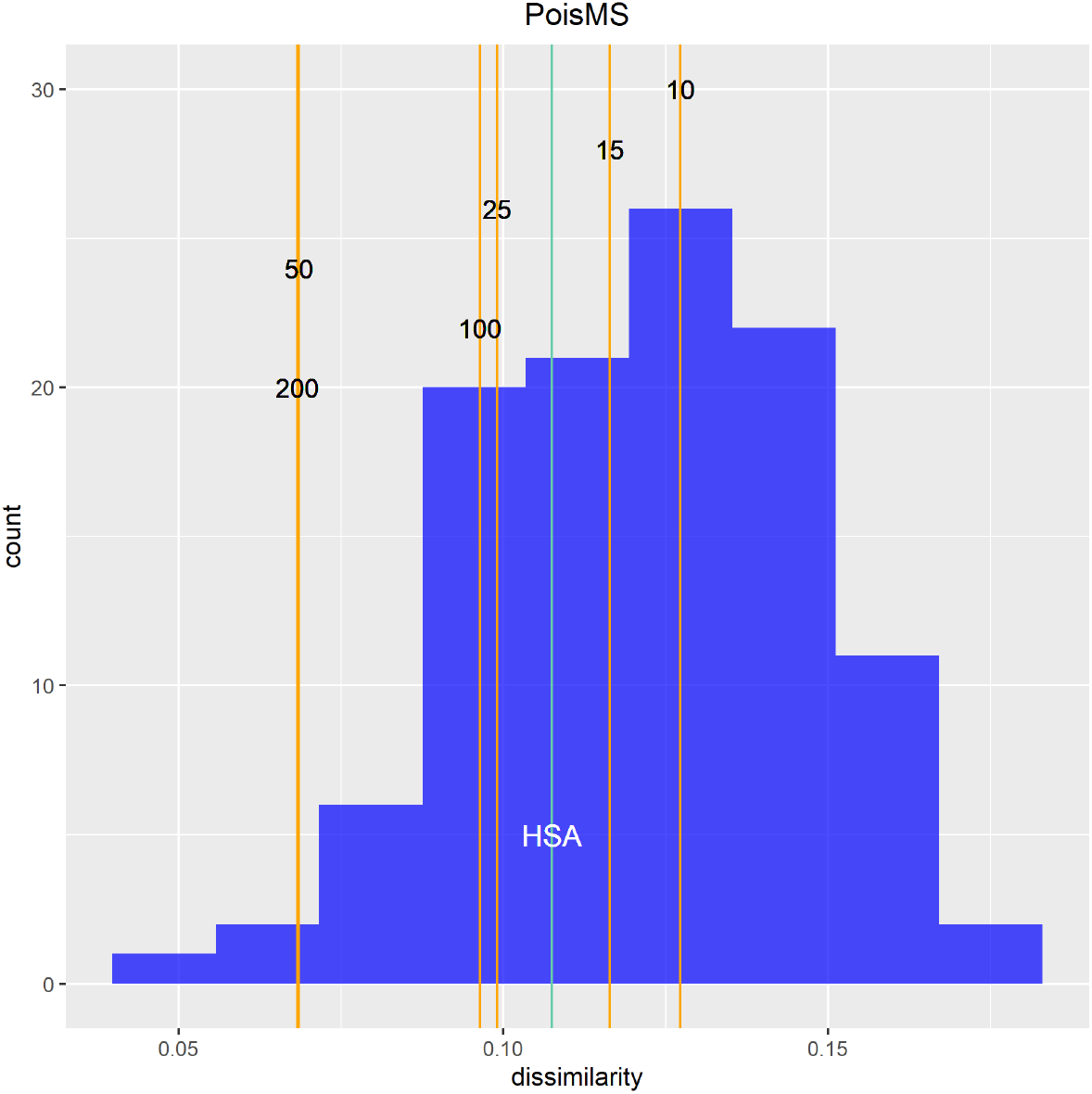
Reference distribution measuring the dissimilarity between the gold standard 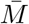 and 111 multiplex FISH replicate conformations *M_i_* for chromosome 20. The vertical orange lines correspond to the dissimilarity between 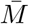 and the low-resolution reconstruction 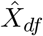 calculated via PoisMS for different *df* values; the light blue line corresponds to the HSA reconstruction (see Sections 2.9 and 2.10).

The following conclusions can be drawn from Figure 4. For chromosome 20 (see the Supplement S10 for chromosome 21), all fits for PoisMS lie within the range of the multiplex FISH dissimilarity distribution that reflects experimental variation. The fact that the PoisMS dissimilarity values are in the left tail of this distribution indicates the accuracy of the proposed reconstructions, highlighting the utility of the proposed methodology. Further, that larger dissimilarity values pertain for HSA, particularly for chromosome 21, suggests that PoisMS performs at least comparably to this well benchmarked alternative. That PoisMS wall clock times are minutes rather than days for HSA is notable.

## 4 Discussion

Central to our principal curve based approaches to 3D chromatin reconstruction is that the configuration of an individual chromosome within the nucleus can be treated as a contiguous 1D curve since the diameter of the chromatin fiber is negligible compared to the nuclear volume. The extent to which the curve is “smooth” is determined by an adaptively selected degrees-of-freedom parameter. As mentioned in the introduction, previous reconstruction methods either impart contiguity indirectly by prescribing constraints, which are difficult to specify, or impose it post hoc. In comparison, our methods based on principal curves are computationally efficient, readily scale to high resolution contact data and are parsimonious with regard tuning parameters.

Our implementation of PoisMS utilizes cubic spline basis functions, which contribute to this computational efficiency. However, the nature of chromatin folding and attendant Hi-C data is such that these bases will be less effective in capturing fine 3D structure, as opposed to global backbone architecture. This derives from the hierarchical, domain-based organization of chromatin, aspects that have been tackled by some reconstruction algorithms using strategies that synthesize solutions obtained at differing scales (Rieber and Mahony, 2017; Trieu *and others*, 2019). We will investigate whether principal curve solutions can similarly serve as building blocks in addition to exploring the use of alternate basis functions, notably wavelets.

Our analyses of Hi-C data from IMR90 cells was motivated by the availability of corresponding multiplex FISH data enabling accuracy assessment. However, the extent and resolution of multiplex FISH imaging is limited, narrowing the applicability of this means of evaluation. An even more fundamental issue pertains to attempting chromatin reconstruction using *bulk* Hi-C data from large cell populations. As has been emphasized (Lando *and others*, 2018), the presence of numerous conflicting contacts suggests that the notion of a consensus underlying 3D conformation is questionable and that there is substantial cell-to-cell structural variation. This places a premium on pursuing single cell reconstructions as enabled by the recent emergence of single cell Hi-C protocols (Ramani *and others*, 2017). That one of these advances (Stevens *and others*, 2017) also provides parallel imaging data, putatively enabling reconstruction accuracy determination, underscores the importance of applying reconstruction methods in single cell settings, despite contact map sparsity, and is the subject of future work.

## 5 Software

Proposed methods are implemented in the R package `PoisMS`; the software is available from Github (https://github.com/ElenaTuzhilina/PoisMS).

## Acknowledgments

The authors thank the Associate Editor and two reviewers from Biostatistics Journal for very helpful comments, including critical appraisal of our original approach, which led to substantial improvements in methodology.

## Supplementary Material

### S1 PCMS lemma proof

#### Lemma 2.1

If 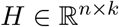 is a matrix with orthogonal columns, i.e. *H^T^ H* = *I*, and *S*(*X*) = *XX^T^* then problem

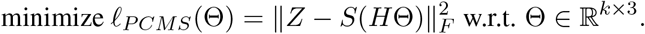

is equivalent to

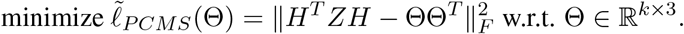

*Proof*. Suppose 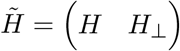 is a column-wise combination of *H* and its orthogonal complement *H*_⊥_. Therefore, 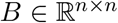 is a square orthogonal matrix and for any 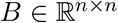 the following relation holds

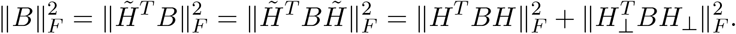

Substituting *B* = *Z* – *H*ΘΘ^*T*^ *H^T^* we conclude that

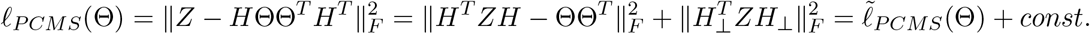

In the last equation the constant term does not depend on the parameter Θ implying the equivalence of the above two optimization problems.

### S2 PCMS reconstruction

In our original development of principal curve metric scaling we *directly* modeled the contact matrix *C* in terms of the inner product matrix *S*(*X*) = *XX^T^*, where 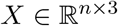 is the matrix of genomic loci coordinates of the reconstruction. Accordingly, in optimizing the *Strain* objective, we are approximating *C_ij_* ≈ 〈*x_i_, x_j_*〉. However, as pointed out by a referee, this is a strong assumption, corresponding to treating *C* as a similarity matrix. The referee further noted that in contrast to *exploratory* uses of MDS, for which postulating plausible similarity matrices suffices, our goal of reproducing (up to a scale factor) the correct 3D chromatin configuration, requires more careful specification of the target similarity matrix.

Here we develop a proposal of the referee in devising one such appropriate similarity matrix, by combining power law transformation with double centering, and demonstrate application of PCMS as a reconstruction tool in this context. Specifically, we first convert contacts to distances via 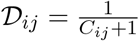. This corresponds to power law transformation with index *α* = −1 (Lesne *and others*, 2014), where we add one in the denominator to avoid dividing by zero. Next, we use the centering operator 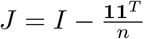 to transform distances to similarities via double centering: 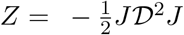 (Buja *and others*, 2008). The heatmap of the resulting similarity matrix is displayed in Figure 5. Finally, we run PCMS on *Z*. The solution 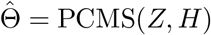 leads to the reconstructions 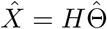 and approximations of the similarity matrix 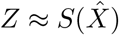 presented in Figure 7 for a range of degrees-of-freedom (*df*) values. The corresponding plot of approximation error

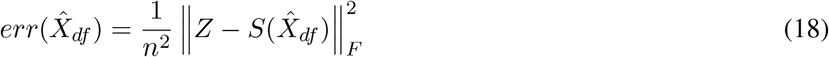

against degrees-of-freedom is presented in Figure 6. Use of segmented regression to identify reasonable model size, given by the elbow, suggests a value *df* = 25 as shown.

**Figure 5:**
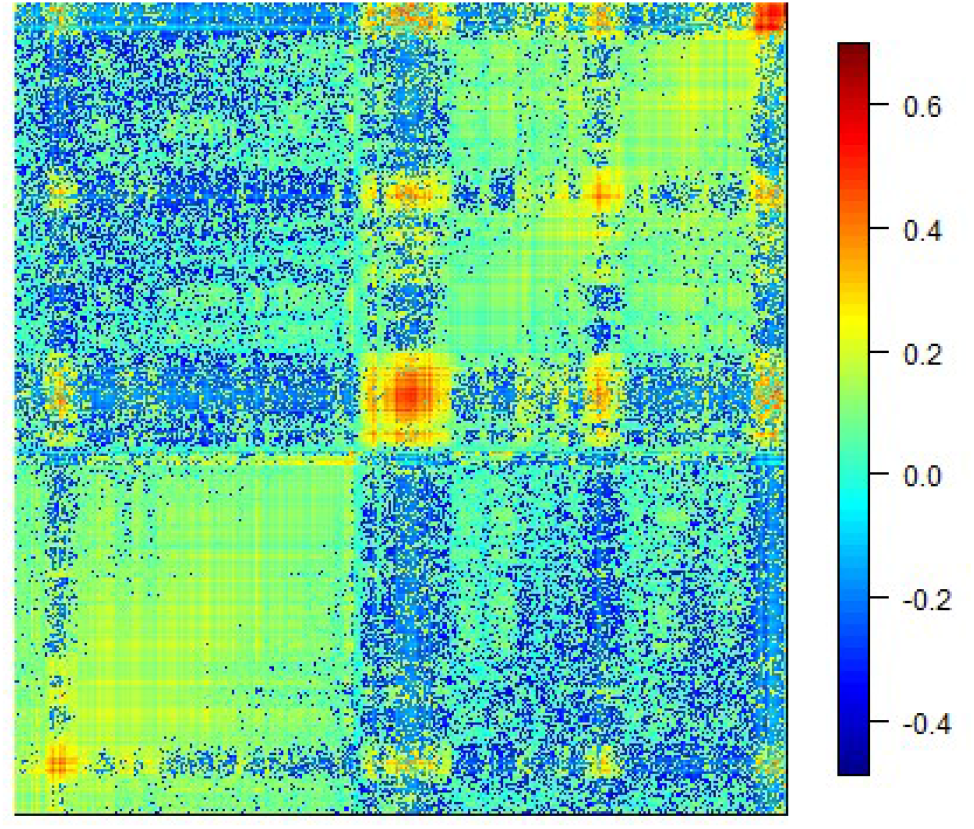
Transformed contact matrix 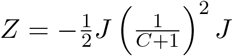.

**Figure 6:**
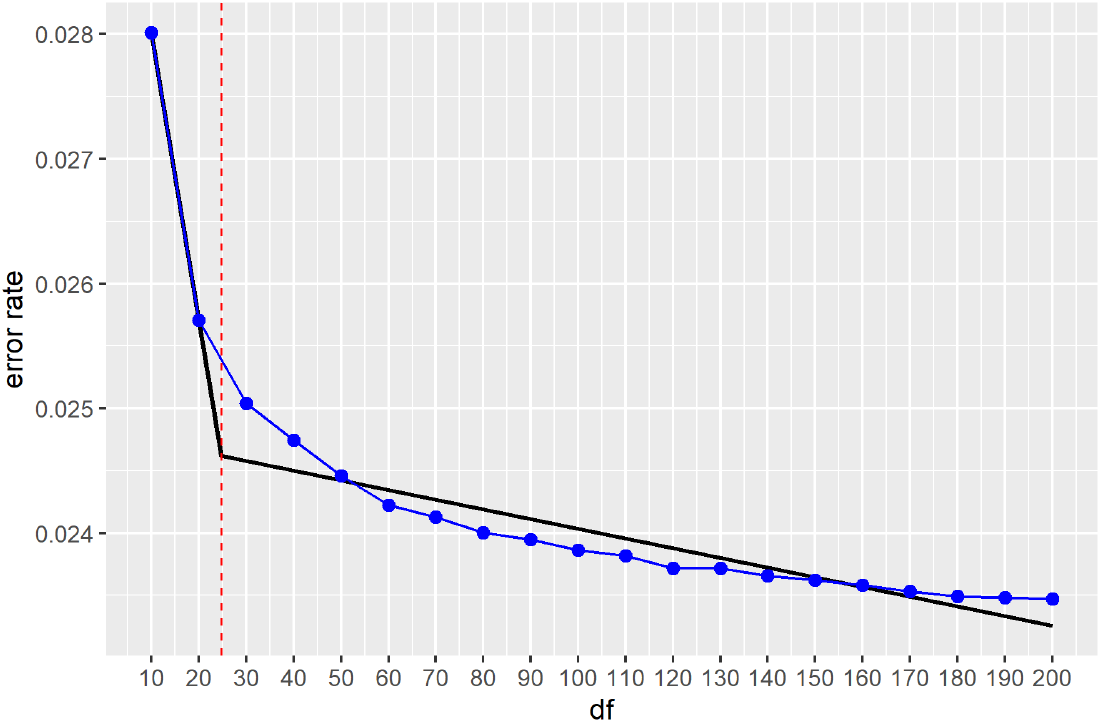
Error rate 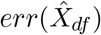 vs. degrees-of-freedom *df* plot for the PCMS approach. The segmented regression is given by the piecewise linear fit (black) with the degrees-of-freedom selected via kink estimation indicated by the red vertical line and segmentation change point corresponding to *df* = 25.

**Figure 7:**
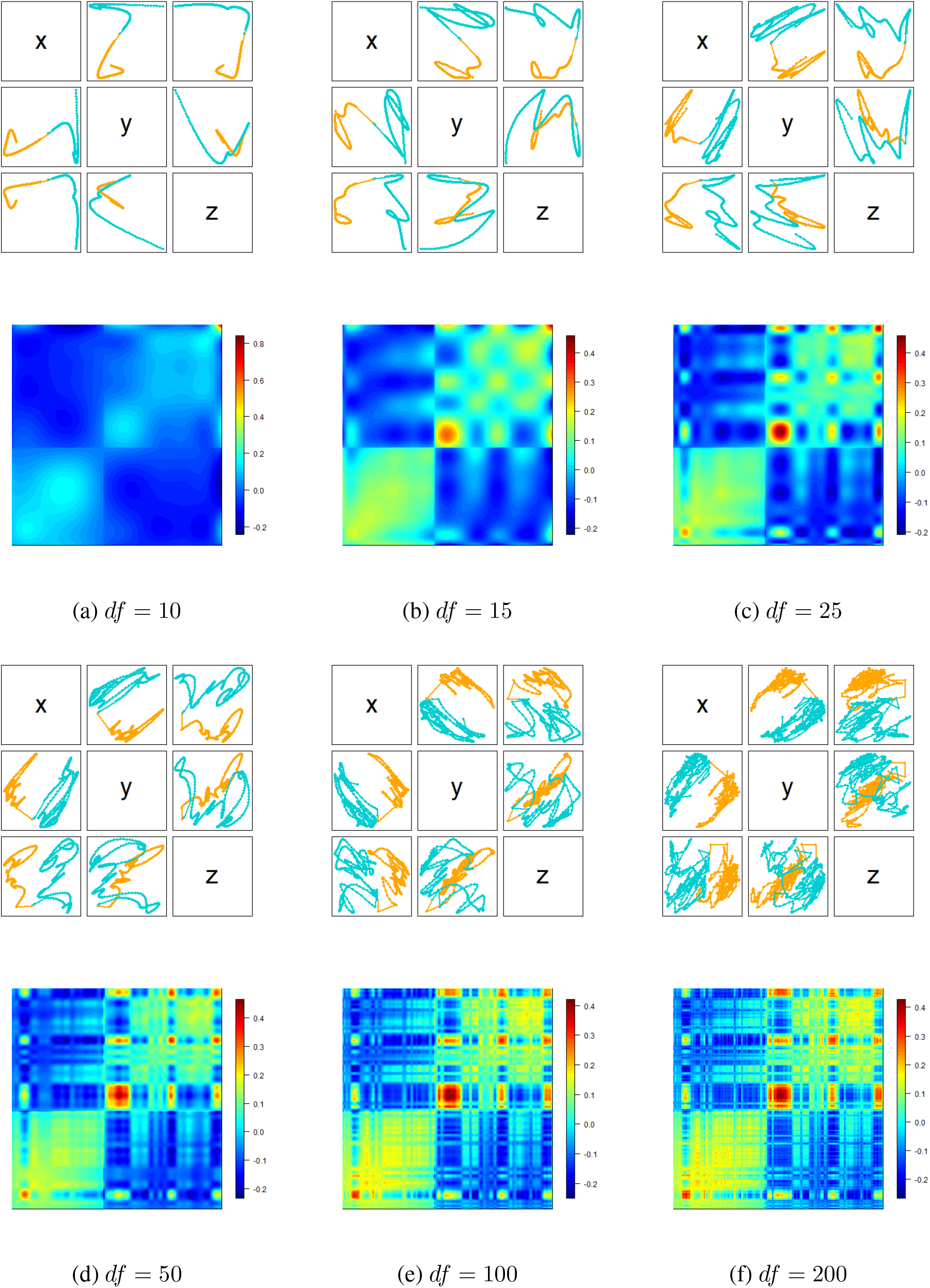
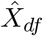, the projections of the resulting reconstruction, with colors (orange, teal) distinguishing chromosome arms, and 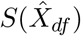, the approximation of *Z*, obtained via PCMS for different degrees-of-freedom values *df*.

### S3 WPCMS lemma proof

#### Lemma 2.2

Let *D*^2^ denote the matrix of squared distances corresponding to the inner product matrix *S* = *XX^T^*. For similarity matrix *Z* and weight matrix *W* let *G* = *W* * (*Z* – *D*^2^) and *G*^+^ = diag(*G* · **1**), a diagonal matrix with column sums of *G* on the diagonal. Then, up to a scaling factor, ∇*ℓ*_*WPCMS*_(*S*) = *G* – *G*^+^.

*Proof*. Note that

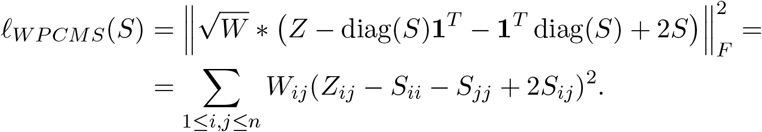

So the partial derivatives are

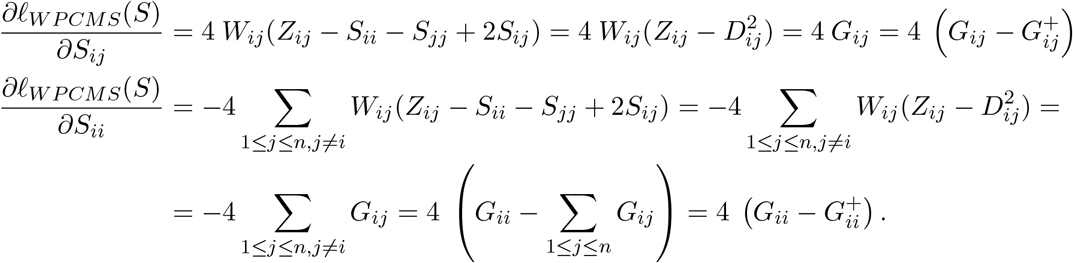

In matrix form this is exactly ∇*ℓ*_*WPCMS*_(*S*) = *G* – *G*^+^.

### S3 WPCMS algorithm extensions

#### S3.1 Learning rate

Since the WPCMS algorithm is based on Projected Gradient Descent (PGD), many extensions can be naturally derived. For example, a learning rate *δ* can be introduced by replacing *G* by *δG* at the **[Gradient]** step. Further, line search can be readily added to the algorithm: **[Gradient]** *S*:= *S* – (*G* – *G*^+^) ⇒

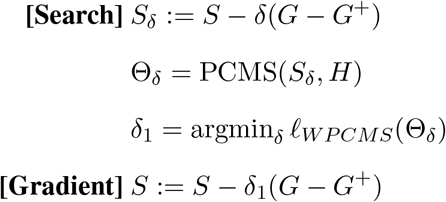

#### S3.2 Intercept

As with PoisMS, it is possible to incorporate an intercept parameter into the WPCMS problem statement, leading to the optimization problem:

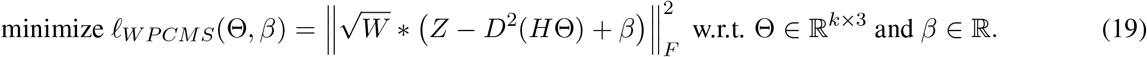

The WPCMS iterative algorithm is then modified by the addition of the following step:

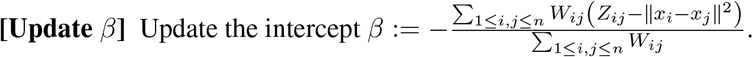

### S4 WPCMS reconstruction

Similarly to PCMS, we can use WPCMS as a standalone chromatin reconstruction technique. To make results comparable to PoisMS we adopt the following approach. According to the Poisson model under consideration, the matrix of Poisson rate parameters Λ depends on genomic loci spatial coordinates *X* via the relation log Λ = −*D*(*X*)^2^ + *β*. Since Λ is the expectation of the contact counts *C* it is reasonable to seek for the approximation

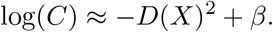

We introduce the binary matrix of weights *W* with elements

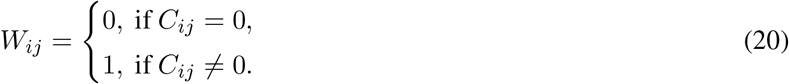

Then, the corresponding optimization problem

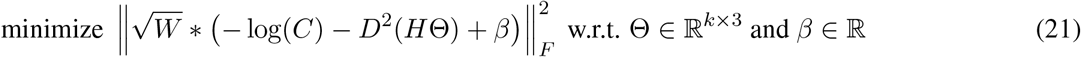

coincides with the WPCMS problem (19) with *Z* = −log(*C*). We obtained the solution 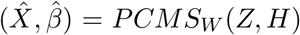 using our WPCMS algorithm and present chromatin reconstructions 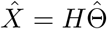 as well as approximations 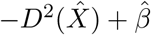 to the log-transformed contact matrix log(*C*) for a series of degrees-of-freedom in Figure 8. The corresponding plot of approximation error

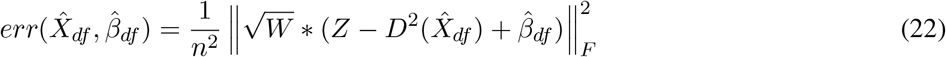

is presented in Figure 9. use of segmented regression to identify reasonable model size, given by the elbow, suggests a value *df* = 35 as shown.

**Figure 8:**
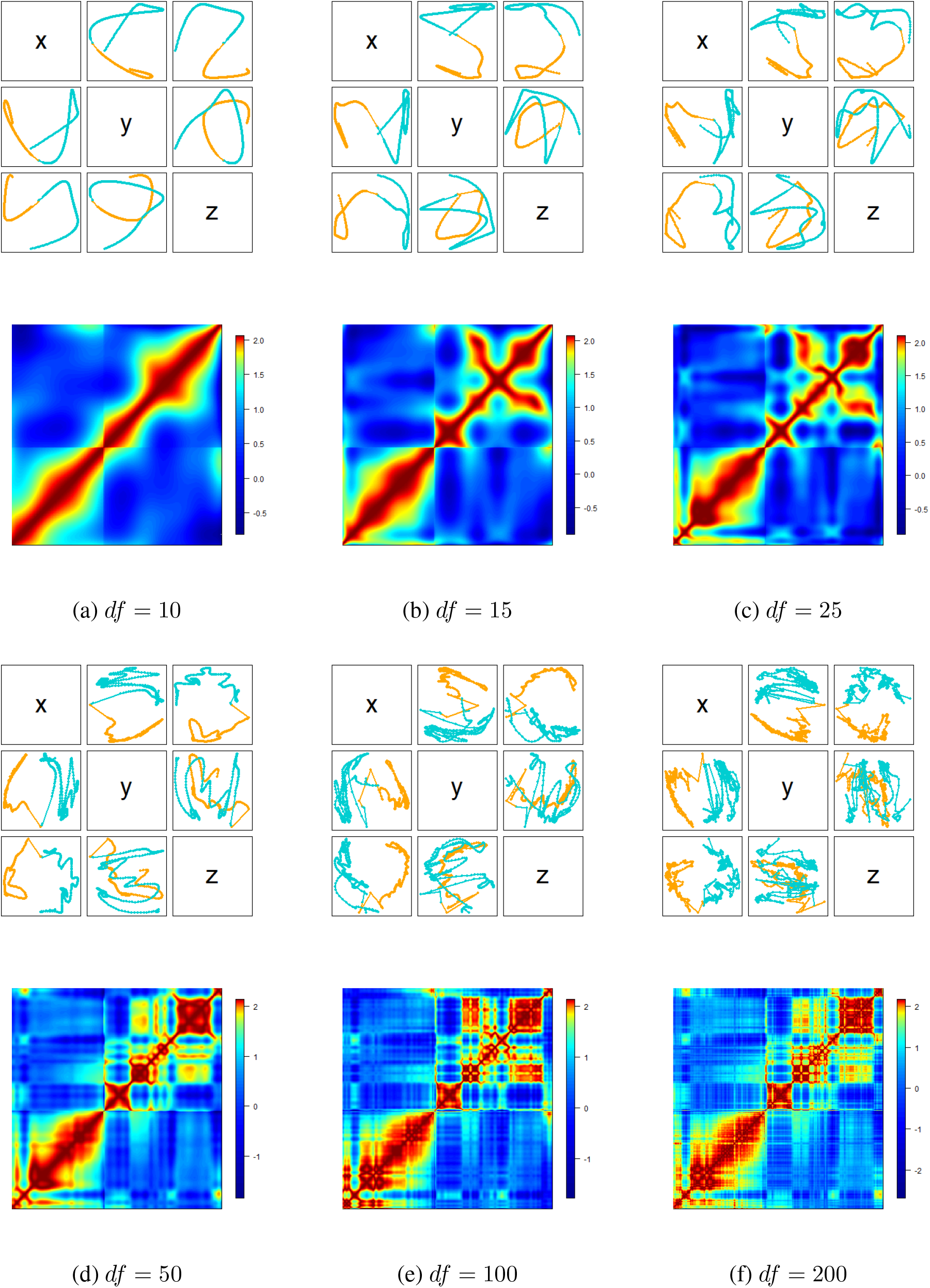
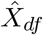, the projections of the resulting reconstruction, with colors (orange, teal) distinguishing chromosome arms, and 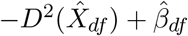, the approximation of log(*C*), obtained via WPCMS for different degrees-of-freedom values *df*.

**Figure 9:**
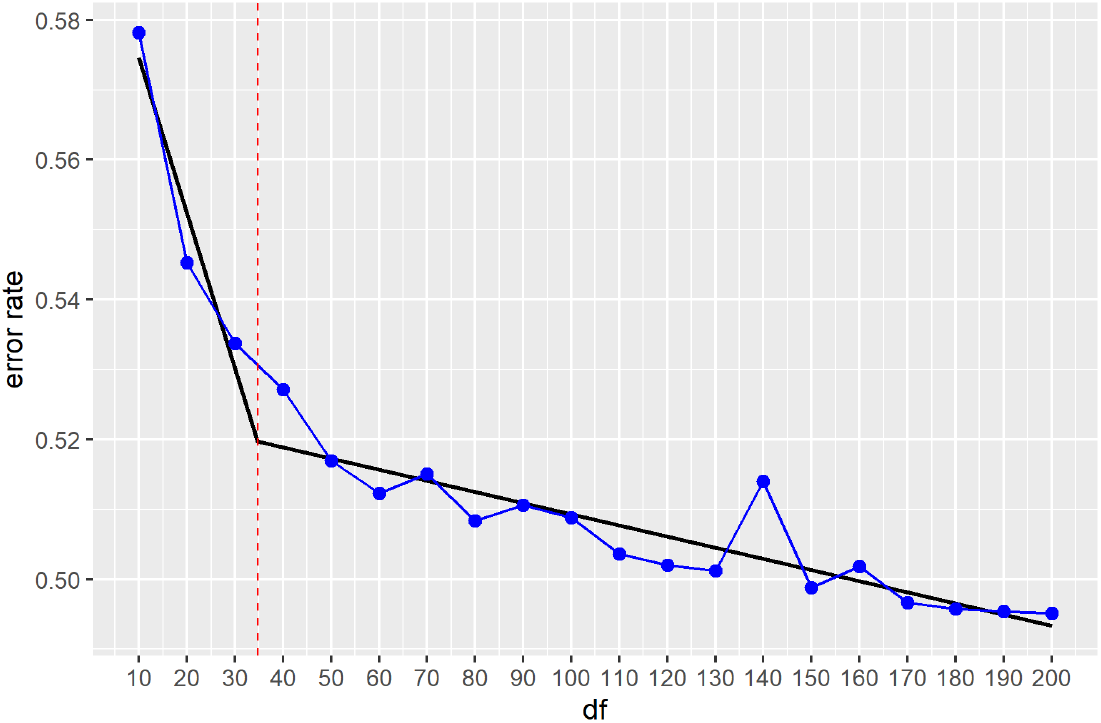
Error rate 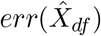 vs. degrees-of-freedom *df* plot for the WPCMS approach. The segmented regression is given by the piecewise linear fit (black) with the degrees-of-freedom selected via kink estimation indicated by the red vertical line and segmentation change point corresponding to *df* = 35.

### S5 WPCMS initialization

We study the influence of the initialization on the resulting WPCMS reconstructions. In our experiments we generated 50 random Θ with 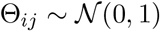. At the **[Initialize]** step for each of these Θ we set the starting value for the reconstruction to *X* = *H*Θ, for fixed degrees-of-freedom *df* = 50, and run the WPCMS algorithm. We then compare the solutions obtained, as well as losses and *β* values. In Figure 10 we superpose coordinate-wise representations of all 50 curves obtained via WPCMS. The WPCMS algorithm converges to two stable conformations. The relative change in the achieved values for both loss and *β* does not exceed 1%; the number of iterations required for the same convergence criterion varies from tens to a few hundreds.

**Figure 10:**
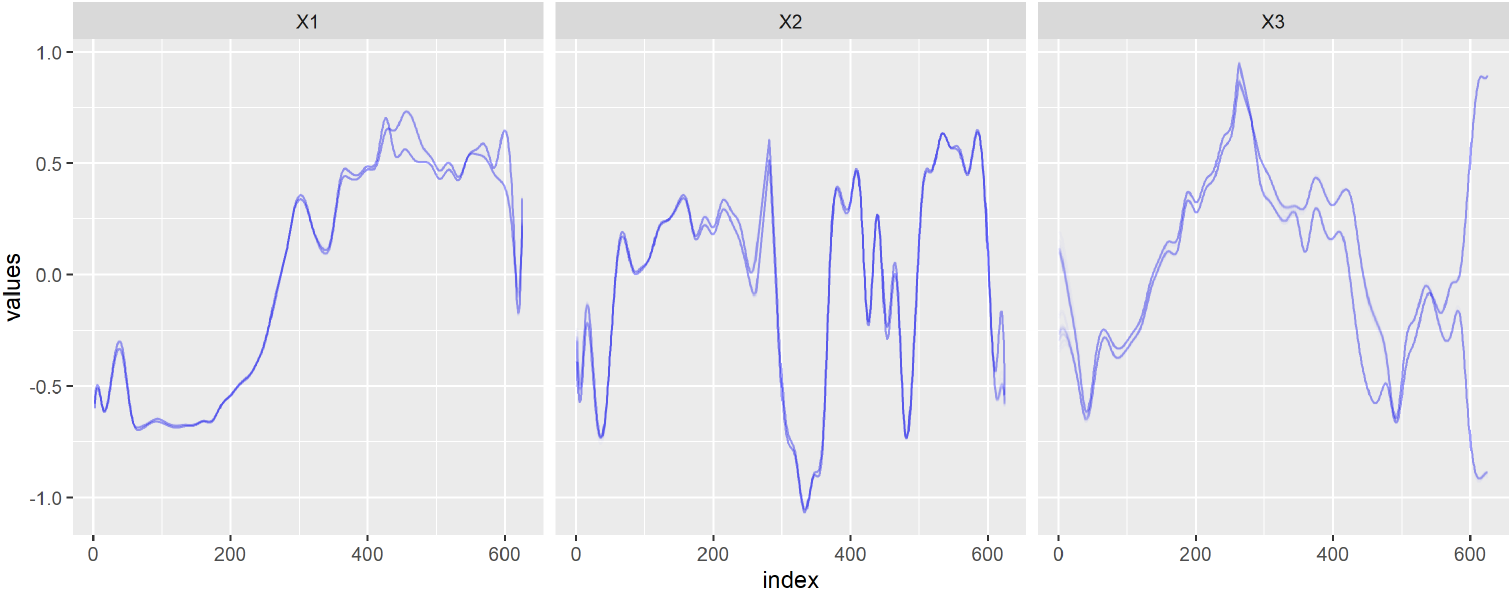
50 reconstructions 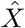 computed for *df* = 50 via WPCMS: each reconstruction corresponds to a random initialization and is represented coordinate-wise. Two largely similar stable conformations are obtained.

Figure 11 depicts the 50 WPCMS solutions in terms of two key geometric characteristics of the reconstructions: distances between two successive genomic loci which reflects curvature, and angles formed by three successive genomic loci which reflects torsion. The plot showcases highly similar geometric structure across reconstructions, irrespective of initialization.

**Figure 11:**
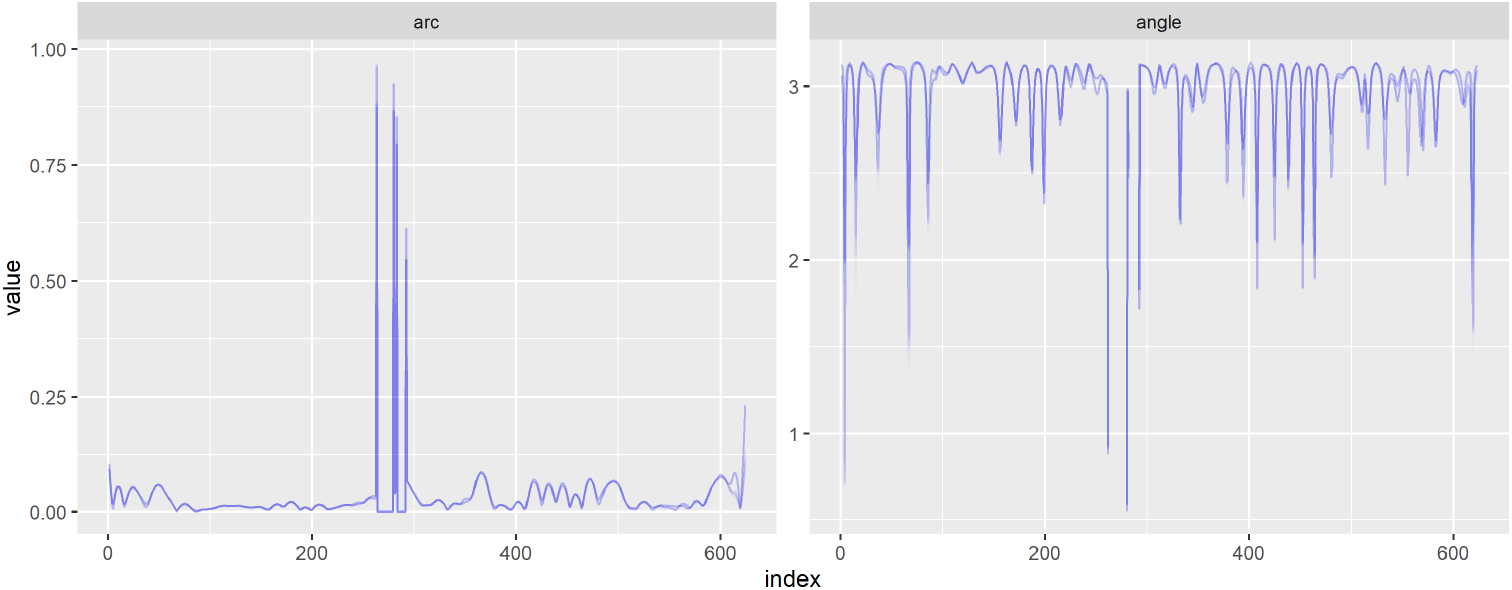
50 reconstructions 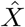 computed for *df* = 50 via WPCMS; each reconstruction corresponds to a random initialization, and is represented as lengths of reconstruction edges and angles between successive edges.

### S6 PoisMS algorithm extensions

#### S6.1 Iterations

It is not necessary to iterate until convergence of the objective *ℓ_SOA_*(Θ) at the **[WPCMS]** step. Based on our experiments, the PoisMS algorithm can be significantly accelerated by performing only a few steps of WPCMS between each update of *W* and *Z*. This can be readily accomplished by varying the accuracy rate *ϵ*_1_ for the WPCMS stopping rule 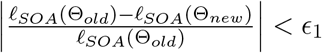.

#### S6.2 Learning rate

Note that the update for the Poisson generalized linear model (GLM) amounts to one step of the Newton method in the space of the natural parameter 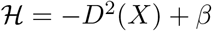. It is not difficult to show that it is equivalent to the Newton step in the parameter space of distances *D*^2^(*X*) and, consequently, in the parameter space of inner products *S*(*X*) since there is a linear dependence of the elements of *D*^2^(*X*) on the elements of *S*(*X*). Therefore, we can add a learning rate as well as line search (in the parameter space of *S*(*X*)) to the PoisMS algorithm as follows:

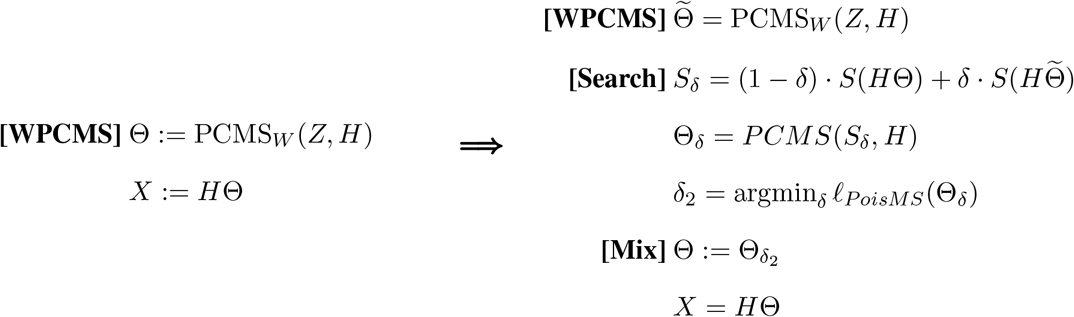

### S7 PoisMS initialization

Akin to our exploration of the impact of initialization on WPCMS (Section S5) here we examine initialization in the context of PoisMS. In Figure 12 we again superpose 50 curves obtained from different initializations of PoisMS and represent them coordinate-wise. As is apparent, PoisMS exhibits higher sensitivity to the starting value of Θ than WPCMS. However, the relative change in the achieved values for both loss and *β* is still small, i.e. does not exceed 3%.

**Figure 12:**
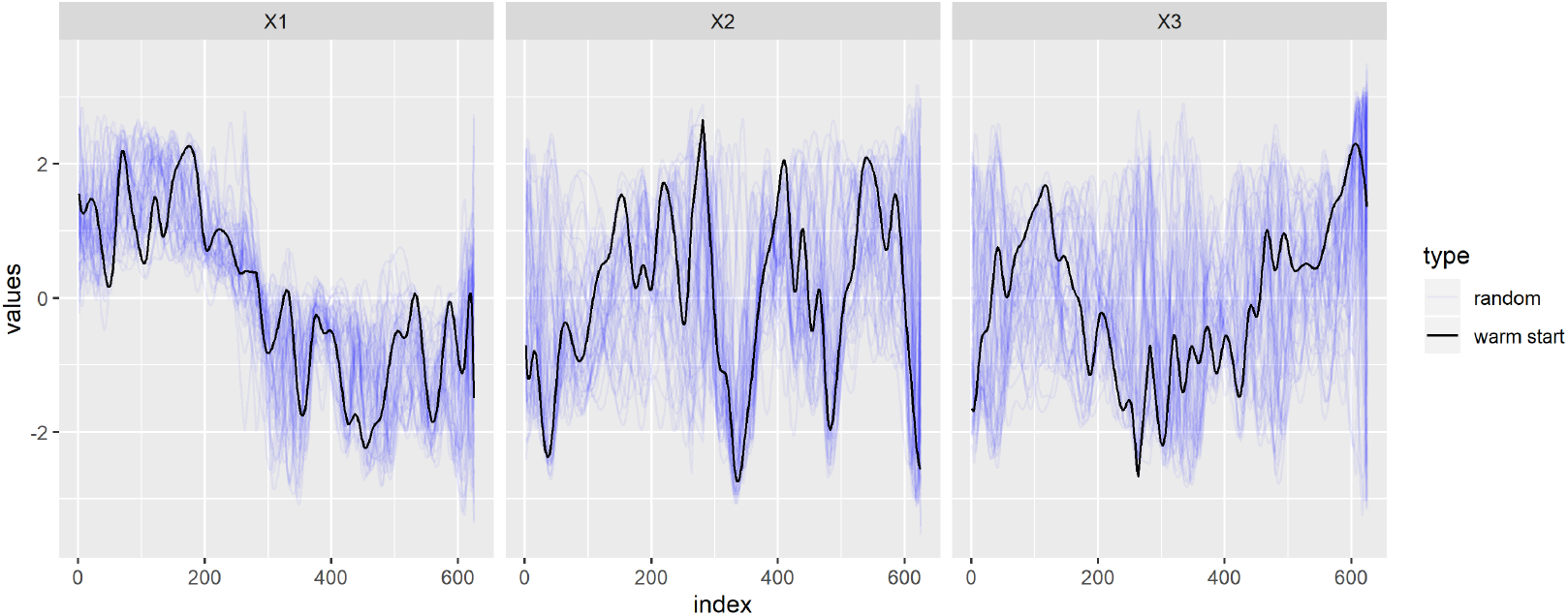
50 reconstructions 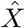 computed for *df* = 50 via PoisMS; each reconstruction corresponds to an initialization (random initialization in blue, warm start in black) and is represented coordinate-wise.

While Figure 12 highlights reconstruction variability with respect to initialization, Figure 13 which, as per Figure 11, captures geometry via successive edge lengths and angles, suggests the existence of a common underlying similar structure.

**Figure 13:**
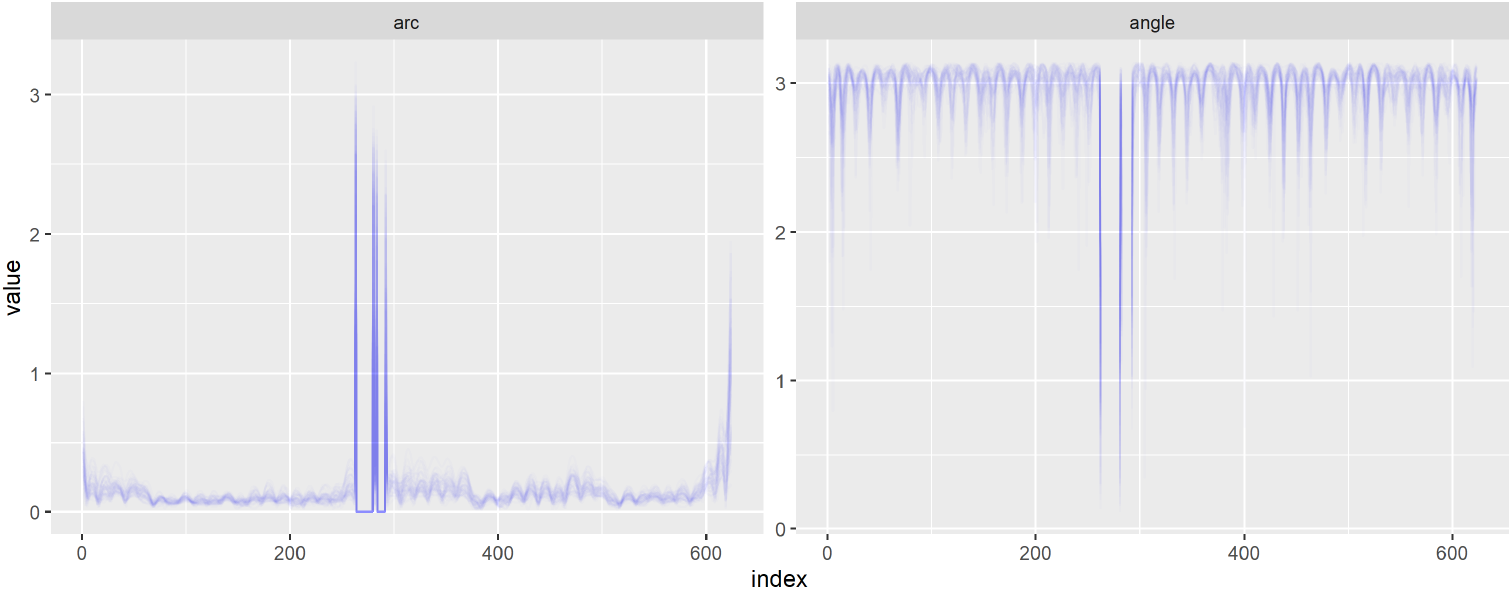
50 reconstructions 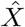 computed for *df* = 50 via PoisMS; each reconstruction corresponds to a random initialization, and is represented as lengths of reconstruction edges and angles between successive edges.

### S8 PoisMS warm start

In view of the sensitivity of PoisMS to initialization we pursue use of a warm start. Based on the heuristic from Section S4, we replace the random initialization step by initializing Θ at the WPCMS solution (21)

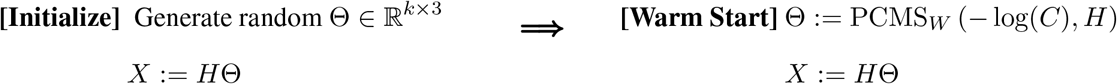

where *W* is the binary matrix (20). Adoption of warm starts significantly decreases the number of iterations required for convergence and allows for lower loss value. The resulting reconstruction is presented in Figure 12 in black.

### S9 Computational complexity

The computational complexity of PCMS is *O*(*n*^2^*k*) operations for computing the product *H^T^ CH* and extra *O*(*k*^3^) for eigen-decomposition (negligible for small *k*). Each iteration of WPCMS has the same complexity as PCMS plus *O*(*n*^2^) operations for computing Hadamard products with matrix *W*. Finally, the complexity of a PoisMS iteration is equivalent to a WPCMS iteration and requires an additional *O*(*n*^2^) operations for calculating the second order approximation.

### S10 PoisMS reconstructions for IMR90 cell chromosome 21

We provide results for IMR90 chromosome 21 at 100kb resolution that parallel the presentation for chromosome 20.

**Figure 14:**
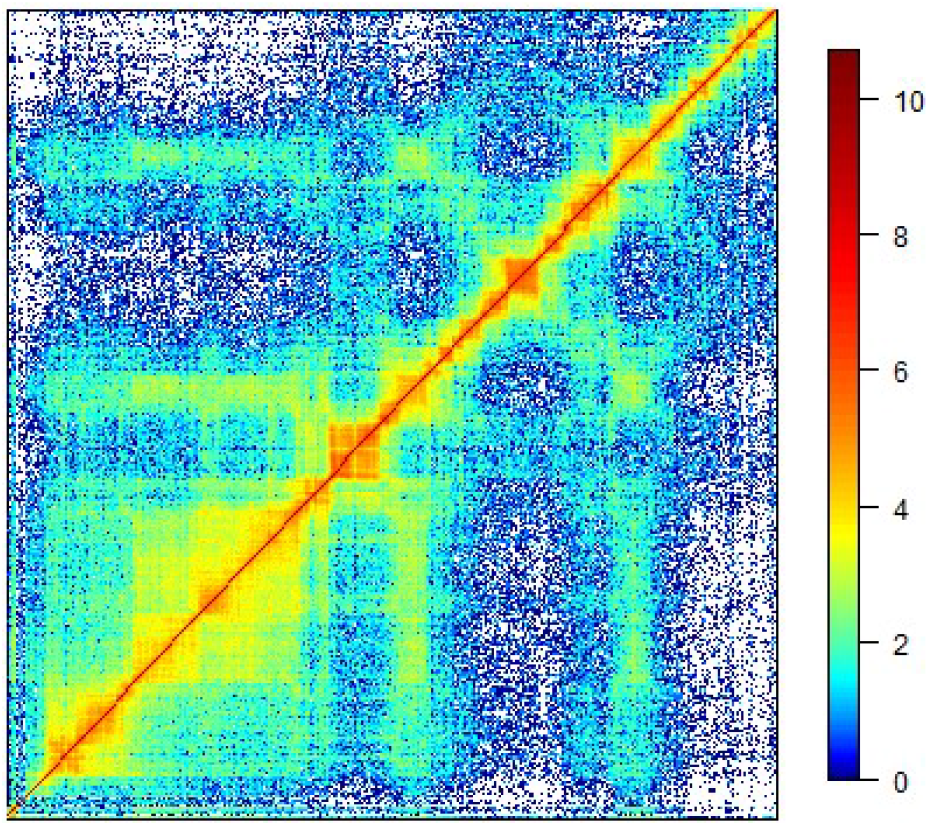
Log-transformed contact matrix log(*C*). White color corresponds to *C_ij_* = 0 or, equivalently, log(*C_ij_*) = −∞.

**Figure 15:**
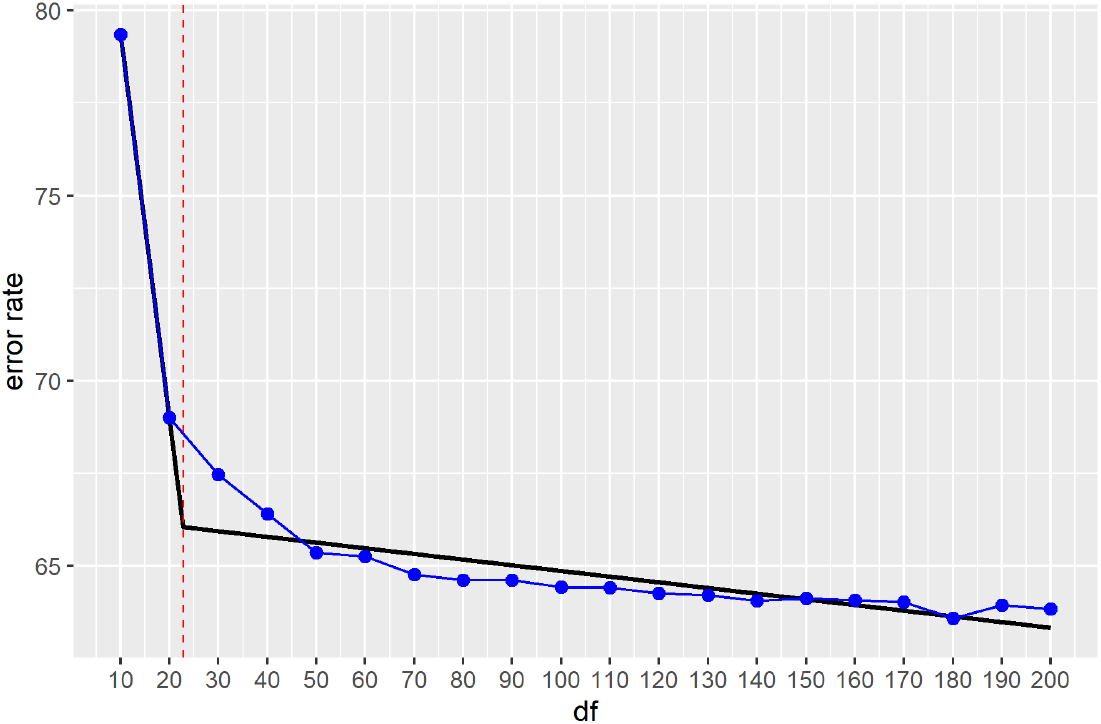
Error rate 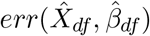 vs. degrees-of-freedom *df* plot for the PoisMS approach. The segmented regression is given by the piecewise linear fit (black) with the degrees-of-freedom selected via kink estimation indicated by the red vertical line and segmentation change point corresponding to *df* = 23.

**Figure 16:**
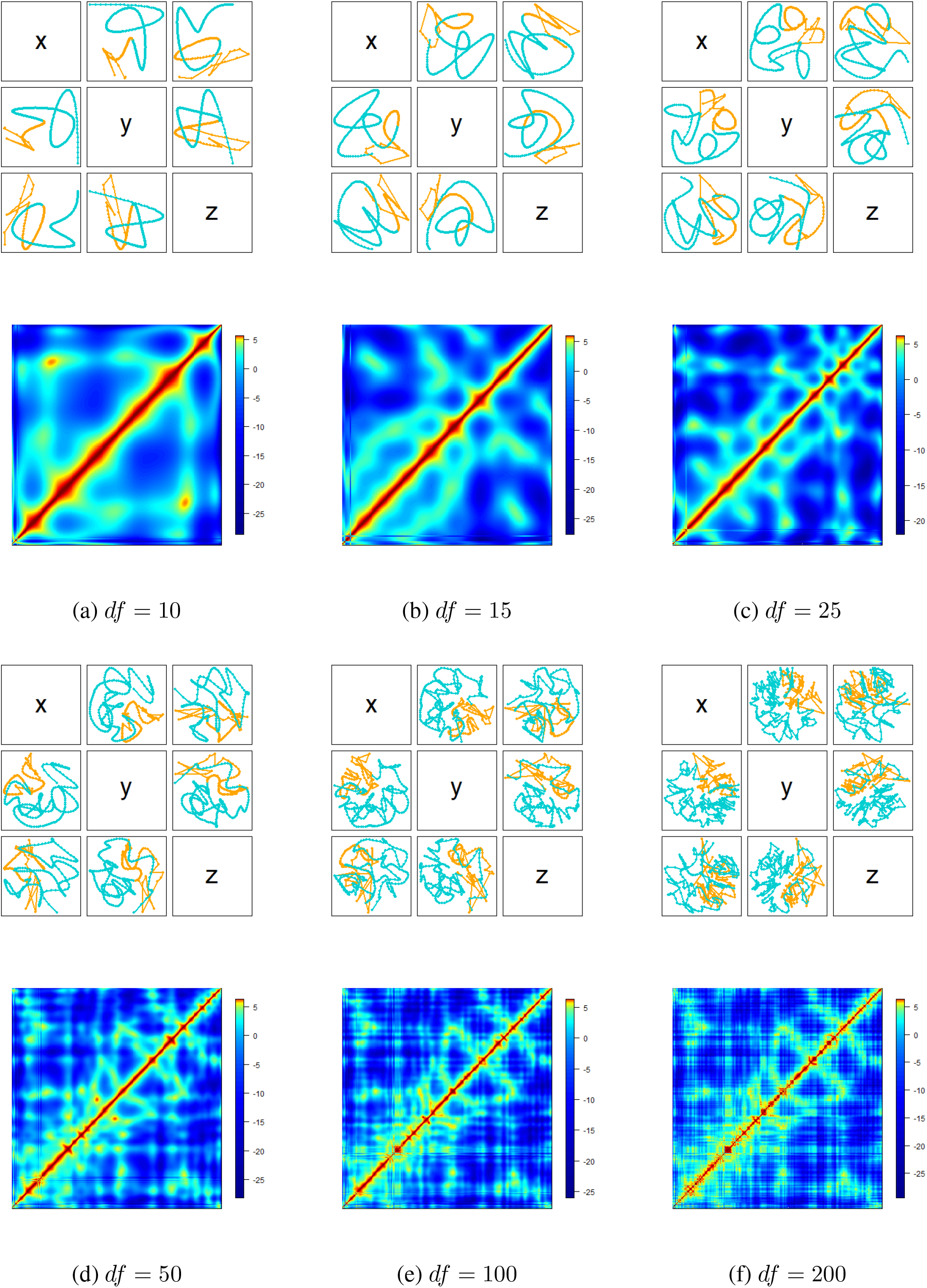
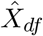, the projections of the resulting reconstruction, with colors (orange, teal) distinguishing chromosome arms, and 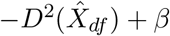, the approximation of log(*C*), obtained via PoisMS for different degrees-of-freedom values *df*.

**Figure 17:**
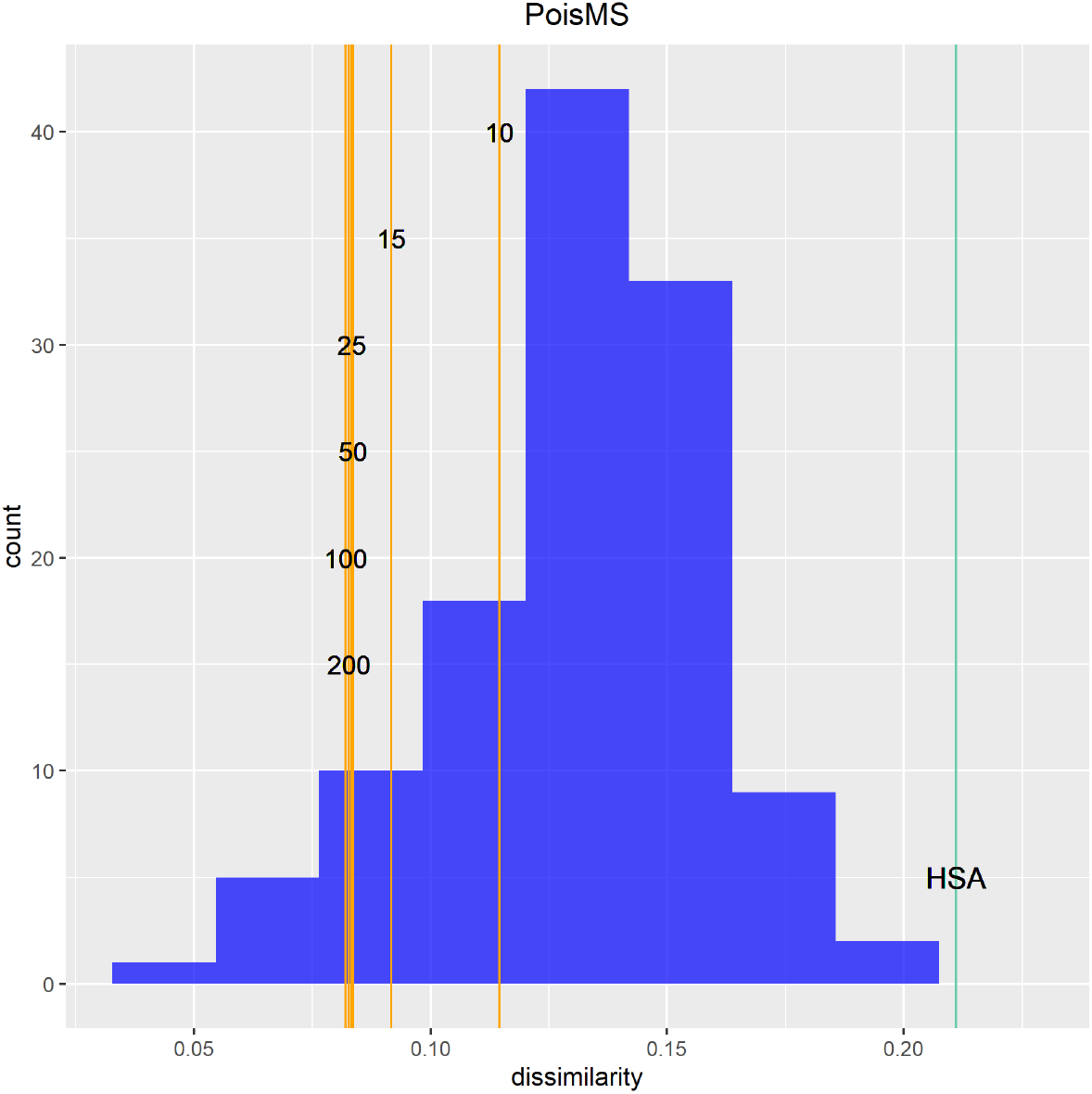
Reference distribution measuring the dissimilarity between the gold standard 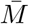 and 111 multiplex FISH replicate conformations *M_i_* for chromosome 21. The vertical orange lines correspond to the dissimilarity between 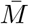 and the low-resolution reconstruction 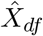 calculated via PoisMS for different *df* values; the light blue line corresponds to the HSA reconstruction.

## References

Ay, F., Bunnik, E. M., Varoquaux, N., Bol, S. M., Prudhomme, J., Vert, J. P., Noble, W. S. and Le Roch, K. G. (2014). Three-dimensional modeling of the P. falciparum genome during the erythrocytic cycle reveals a strong connection between genome architecture and gene expression. Genome Research 24, 974–88.

Breiman, L., Friedman, J. H., Olshen, R. A. and Stone, C. J. (1984). Classification and Regression Trees. New York: Chapman and Hall.

Buja, A., Swayne, D. F., Littman, M. L., Dean, N., Hofman, H. and Chen, L. (2008). Data Visualization With Multidimensional Scaling. Journal of Computational and Graphical Statistics 17, 444–472.

Capurso, D., Bengtsson, H. and Segal, M. R. (2016). Discovering hotspots in functional genomic data superposed on 3D chromatin configuration reconstructions. Nucleic Acids Research 44, 2028–2035.

Capurso, D. and Segal, M. R. (2014). Distance-based assessment of the localization of functional annotations in 3D genome reconstructions. BMC Genomics 15, 992.

Caudai, C., Salerno, E., Zopp, M. and Tonazzini, A. (2015). Inferring 3d chromatin structure using a multiscale approach based on quaternions. BMC Bioinformatics 16, 234.

Dekker, J., Rippe, K., Dekker, M. and Kleckner, N. (2002). Capturing chromosome conformation. Science 295, 1306–1311.

Dixon, J. R., Selvaraj, S., Yue, F., Kim, A., Li, Y., Shen, Y., Hu, M., Liu, J. S. and Ren, B. (2012). Topological domains in mammalian genomes identified by analysis of chromatin contacts. Nature 485, 376–380.

Duan, Z., Andronescu, M., Schutz, K., McIlwain, S., Kim, Y. J., Lee, C., Shendure, J., Fields, S., Blau, C. A. and Noble, W. S. (2010). A three-dimensional model of the yeast genome. Nature 465, 363–367.

Fudenberg, G. and Mirny, L. A. (2012). Higher-order chromatin structure: bridging physics and biology. Current Opinions in Genetics & Development 22, 115–124.

Hastie, T. J. and Stuetzle, W. (1989). Principal curves. Journal of the American Statistical Association 406, 502–516.

Hastie, T. J., Tibshirani, R. J. and Friedman, J. H. (2009). The Elements of Statistical Learning. New York: Springer.

Hastie, T. J., Tibshirani, R. J. and Wainwright, M. J. (2015). Statistical Learning with Sparsity: The Lasso and Generalizations. New York: Chapman and Hall.

Hutchins, L. N., Murphy, S. M., Singh, P. and Graber, J. H. (2008). Position-dependent motif characterization using non-negative matrix factorization. Bioinformatics 24, 2684–2690.

Jolliffe, I. (2002). Principal Component Analysis. New York: Springer.

Kruskal, J. B. and Wish, M. (1978). Multidimensional Scaling. Newbury Park: Sage.

Lando, D., Stevens, T. J., Basu, S. and Laue, E. D. (2018). Calculation of 3D genome structures for comparison of chromosome conformation capture experiments with microscopy: An evaluation of single-cell Hi-C protocols. Nucleus 9, 190–201.

Lee, C. S., Wang, R. W., Chang, H. H., Capurso, D., Segal, M. R. and Haber, J. E. (2016). Chromosome position determines the success of double-strand break repair. Proceedings of the National Academy of Science 113, 146–154.

Lesne, A., Riposo, J., Roger, P., Cournac, A. and Mozziconacci, J. (2014). 3D genome reconstruction from chromosomal contacts. Nature Methods 11, 1141–1143.

Lieberman-Aiden, E., van Berkum, N. L., Williams, L., Imakaev, M., Ragoczy, T., Telling, A., Amit, I., Lajoie, B. R., Sabo, P. J., Dorschner, M. O., Sandstrom, R., Bernstein, B., Bender, M. A., Groudine, M., Gnirke, A., Stamatoyannopoulos, J., Mirny, L. A., Lander, E. S. and others. (2009). Comprehensive mapping of long-range contacts reveals folding principles of the human genome. Science 326, 289–293.

Mitelman, F., Johansson, B. and Mertens, F. (2007). The impact of translocations and gene fusions on cancer causation. Nature Reviews Cancer 7, 233–245.

Muggeo, V. M. (2008). segmented: an R package to fit regression models with broken-line relationships. Rnews 8, 20–25.

Oksanen, J., Blanchet, F. G., Friendly, M., Kindt, R., Legendre, P., McGlinn, D., Minchin, P. R., O’Hara, R. B., Simpson, G. L., Solymos, P., Stevens, H., Szoecs, E. *and others.* (2016). vegan: Community Ecology Package. R package version 2, 4–1.

Park, J. and Lin, S. (2017). A random effect model for reconstruction of spatial chromatin structure. Biometrics 73, 52–62.

Ramani, V., Deng, X., Gunderson, K. L., Steemers, F. J., Disteche, C. M., Noble, W. S., Duan, Z. and Shendure, J. (2017). Massively multiplex single-cell Hi-C. Nature Methods 14, 263–266.

Rao, S. S., Huntley, M. H., Durand, N. C., Stamenova, E. K., Bochkov, I. D., Robinson, J. T., Sanborn, A. L., Machol, I., Omer, A. D., Lander, E. S. *and others.* (2014). A 3D map of the human genome at kilobase resolution reveals principles of chromatin looping. Cell 159, 1665–1680.

Rieber, L. and Mahony, S. (2017). miniMDS: 3D structural inference from high-resolution hi-c data. Bioinformatics 33, 261–266.

Rosenthal, M., Bryner, D., Huffer, F., Evans, S., Srivastava, A. and Neretti, N. (2019). Bayesian Estimation of 3D Chromosomal Structure from Single Cell Hi-C Data. Journal of Computational Biology 26, 1191–1202.

Segal, M. R. and Bengtsson, H. L. (2015). Reconstruction of 3D genome architecture via a two-stage algorithm. BMC Bioinformatics 16, 373.

Segal, M. R. and Bengtsson, H. L. (2018). Improved accuracy assessment for 3D genome reconstructions. BMC Bioinformatics 19, 196.

Shavit, Y., Hamey, F. K. and Lio, P. (2014). FisHiCal: an R package for iterative FISH-based calibration of Hi-C data. Bioinformatics 30, 3120–3122.

Stevens, T. J., Lando, D., Basu, S., Atkinson, L. P., Cao, Y., Lee, S. F., Leeb, M., Wohlfahrt, K. J., Boucher, W., O’Shaughnessy-Kirwan, A., Cramard, J., Faure, A. J., Ralser, M., Blanco, E., Morey, L., Sanso, M., Palayret, M. G. S., Lehner, B., Di Croce, L., Wutz, A., Hendrich, B., Klenerman, D. and others. (2017). 3D structures of individual mammalian genomes studied by single-cell Hi-C. Nature 544, 59–64.

Trieu, T., Oluwadare, O. and Cheng, J. (2019). Hierarchical reconstruction of high-resolution 3D models of large chromosomes. Scientific Reports 9, 4971.

Varoquaux, N., Ay, F., Noble, W. S. and Vert, J. P. (2014). A statistical approach for inferring the 3D structure of the genome. Bioinformatics 30, 26–33.

Wang, S., Su, J.-H., Beliveau, B. J., Bintu, B., Moffitt, J. R., Wu, C.-T. and Zhuang, X. (2016). Spatial organization of chromatin domains and compartments in single chromosomes. Science 353, 598–602.

Witten, D. M. and Noble, W. S. (2012). On the assessment of statistical significance of three-dimensional colocalization of sets of genomic elements. Nucleic Acids Research 40, 3849–3855.

Yang, T., Zhang, F., Yardimci, G. G., Song, F., Hardison, R. C., Noble, W. S., Yue, F. and Li, Q. (2017). HiCRep: assessing the reproducibility of Hi-C data using a stratum-adjusted correlation coefficient. Genome Research 27, 1939–1949.

Zhang, Z., Li, G., K.-C., TOH and Sung, W.-K. (2013). 3D chromosome modeling with semi-definite programming and Hi-C data. Journal of Computational Biology 20, 831–846.

Zou, C., Zhang, Y. and Ouyang, Z. (2016). HSA: integrating multi-track Hi-C data for genome-scale reconstruction of 3D chromatin structure. Genome Biology 17, 40.

